# Repurposing of the assembly chaperone PAC1-PAC2 defines a specialized proteasome landscape in mature mammalian sperm

**DOI:** 10.64898/2026.05.31.728944

**Authors:** Xuehai Zhou, Linlin Shen, Xuyang Yuan, Chen Chen, Tingfeng Wang, Zuyang Li, Jiahao Peng, Xianze Lang, Xingyan Ye, Tian Chen, Kaijian Chen, Chen Su, Zhong Huang, Yao Cong

**Author notes:** These authors contributed equally (Xuehai Zhou, Linlin Shen, Xuyang Yuan, Chen Chen, Tingfeng Wang). Correspondence (Y.C.), (Z.H.).

## Abstract

Mature mammalian sperm rely on active proteasomes containing the testis-specific subunit α4s for capacitation, acrosomal exocytosis and fertilization, but how these proteasomes are structurally specialized remains unclear. We determine the endogenous proteasome landscape of mature bovine spermatozoa and testis using cryo-electron microscopy. We resolve five sperm proteasome assemblies: PA200-20S, free 20S, a previously unrecognized PA200-20S-PAC1/2 hybrid, and PAC1/2-20S complexes capped at one or both ends. Unexpectedly, the PAC1/2-bound complexes are mature and POMP-free, selectively incorporated for α4s, lack α2, and adopt a PAC1/2 orientation and gate configuration distinct from known assembly intermediates. Together with elevated proteolytic activity, these features indicate that PAC1/2 is repurposed from a transient assembly chaperone into a stable post-assembly regulator in mature sperm. α4s-containing proteasomes assemble through the canonical PAC1-PAC4/POMP pathway yet may follow β2-β3-β1 incorporation ordering; functionally, α4s incorporation remodels the sCP surface and enhances peptidase activities, definining the spermatoproteasomes as a hyperactive 20S variant. PAC1/2-bound proteasomes are absent from testis, where PA200-20S and 26S proteasomes are present, while 26S proteasomes are nearly lost in mature sperm. Our findings define a sperm-specific, ATP-independent proteasome landscape, expand the functional repertoires of α4s and PAC1/2, and provide a structural framework for proteasome specialization in male fertility.

## Introduction

The ubiquitin-proteasome system is the major proteolytic engine of eukaryotic cells, mediating the selective protein degradation that underlies homeostasis, signaling, and development^1–3^. Its catalytic centre is the 20S core particle (CP), a barrel-shaped protease gated at both ends by heptameric α-subunit rings. Substrate access to this catalytic chamber is controlled by distinct activators that dock onto the α-ring and remodel the gate. The ATP-dependent 19S regulatory particle (RP) assebmels with the 20S to form the 26S proteasome for ubiquitin-dependent degradation^4–8^, whereas ATP-independent activators, including PA200 (PSME4)^9–12^ and PA28^13–15^, drive ubiquitin-independent degradation of specialized substrates^11,16–18^. PA200 is of particular important in the male germline: it is enriched in testis, essential for fertility, and mediates acetylation-dependent histone degradation during spermiogenesis^9–12^.

Spermatogenesis imposes extraordinary, stage-specific proteolytic demands. Meiotic progression, histone-to-protamine exchange, acrosome biogenesis, flagellum assembly, and residual-cytoplasm removal each relies on the timely, selective degradation of distinct protein cohorts^19,20^. Male germ cells have therefore evolved proteasome adaptations that are absent from somatic tissues^21–23^. A central adaptation of the male germline proteasome is the replacement of the constitutive α4 subunit (PSMA7) by its testis-specific paralog α4s (PSMA8), forming the spermatoproteasome (α4s-containing proteasomes)^18,24,25^. Genetic ablation of α4s causes meiotic arrest and male infertility in mice^18,25,26^, and α4s has been proposed to favour PA200 engagement^25,27^, generating PA200-s20S complexes implicated in acetylated histone removal during spermiogenesis^11,23,25,26^. However, how α4s reshapes proteasome architecture, gate dynamics, regulator preference and catalytic activity, and whether α4s-containing proteasomes rely on alternative biogenesis mechanisms remains to be explored.

20S biogenesis is guided by dedicated assembly chaperones^28–31^. In the canonical pathway, the PAC1-PAC2 and PAC3-PAC4 heterodimers scaffold stepwise α-ring assembly and subsequent β-subunit incorporation^32–35^. Previous studies show that PAC1-PAC4 first engages an α4/α5/α6/α7 intermediate, after which αl or α3 is added, with α2 incorporation restricted after α1^36,37^. Both PAC1 and PAC2 contain C-terminal HbYX motifs, shared with PA200 and 19S Rpt subunits, which insert into α-ring pockets and can promote gate opening^38–41^. In canonical assembly, however, the release of PAC1-PAC2 is coupled to the autocatalytic degradation of the maturation factor POMP within the nascent 20S chamber, a checkpoint that marks proteasome maturation^35,36,42^. PAC1-PAC2 has therefore been regarded as a strictly transient assembly factors, absent from mature proteasomes in all systems examined to date^32,33,35,36,42–45^. Whether this rule holds in specialized germ-cell proteasomes remain unclear.

Despite the biological importance of sperm proteolysis, the native proteasome repertoire and its regulation in terminally differentiated spermatozoa remain poorly defined. Recent cryo-electron tomography (cryo-ET) studies have identified 20S^46^, or both 20S and PA200-20S^47^, as the dominant nuclear proteasomes in human sperm lacunae, but these analyses leave open whether additional proteasome states exist in mature sperm or testicular germ cells. Mature spermatozoa are largely transcriptionally and translationally silent, although local translation-coupled protein folding has recently been linked to sustain mammalian sperm flagellar motility^48^. Nevertheless, sperm proteasomes remain enzymatically active and are essential for capacitation, acrosomal exocytosis, and zona pellucida penetration during fertilization^21,49,50^. What proteasome assemblies are present in mature sperm, how they are structurally configured, and how their activity is maintained remain open questions.

Here, we define the endogenous proteasome landscape of bovine mature spermatozoa and testis by cryo-EM. We resolve five sperm proteasome assemblies at 2.76-3.18 Å resolution: PA200-20S, free 20S, a previously unrecognized PA200-20S-PAC1/2 hybrid, and PAC1/2-20S complexes capped at one or both ends. The PAC1/2-bound complexes are mature and POMP-free, selectively incorporate α4s, lack α2, and adopt a PAC1/2 orientation and gate configuration distinct from any known assembly intermediates, indicating that PAC1/2 has been repurposed as a stable post-assembly regulator in mature sperm. Reconstitution in HEK293F cells shows that α2 exclusion is imposed by the mature-sperm cellular environment, not by α4s or PAC1/PAC2 alone. In addition, 26S proteasomes are nearly absent from mature bovine spermatozoa and present in testis, whereas α4s-containing proteasomes assemble using the canonical chaperone machinery yet proceed through a slightly distinct pathway, and are intrinsically hyperactive. Together, these findings expand the functional repertoire of both α4s and PAC1/2, reveal a sperm-tailored, ATP-independent proteasome landscape, and provide a structural framework for understanding proteasome specialization in male fertility.

## Results

### An unexpectedly diverse proteasome landscape in mature bovine spermatozoa

To define the native proteasome repertoire of mature mammalian sperm, we purified endogenous proteasomes from bovine cauda epididymal spermatozoa by ion-exchange chromatography followed by glycerol-gradient centrifugation (Fig. S1A). SDS-PAGE and negative-stain electron microscopy (NS-EM) confirmed sample quality (Fig. S1B-D). Mass spectrometry (MS) identified PA200 together with all 20S CP subunits, including both α4 (PSMA7) and the testis-specific α4s (PSMA8) (Fig. 1A). Immunofluorescence assay revealed that PA200 was mainly localized in the nuclear region of the sperm head and along the entire flagellum (Fig. 1B), consistent with previous reports^9^.

**Fig. 1.**
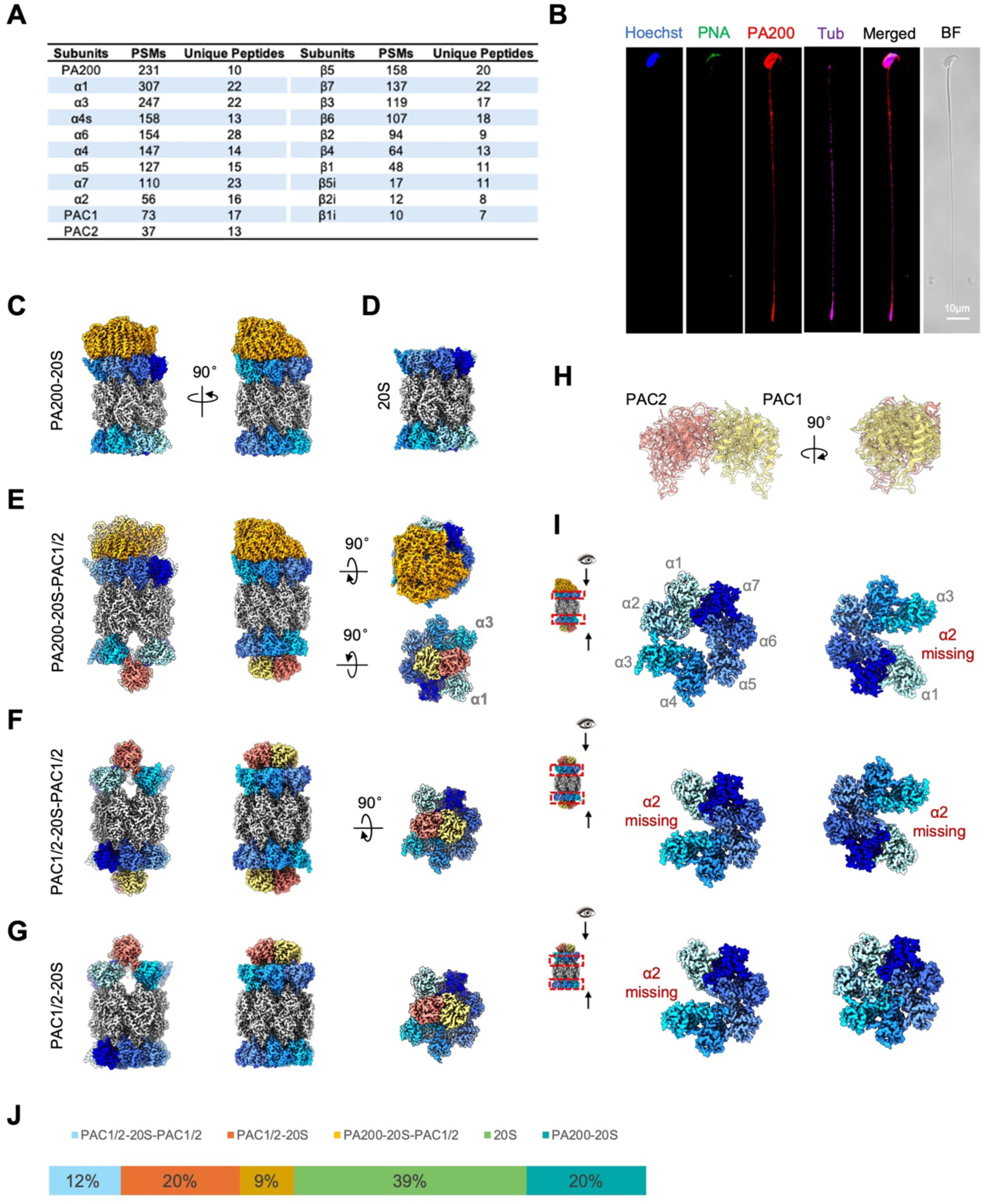
Characterization of mature sperm proteasome complexes. (A) Mass spectrometry analysis of proteasomes purified from mature bovine spermatozoa. (B) Immunofluorescence of mature mouse spermatozoa showing the subcellular localization of PA200. Hoechst (Hoechst33342) labels the nucleus; PNA marks the acrosome; Tub, α-tubulin; BF, bright field. Scale bar: 10μm. (C-G) Cryo-EM reconstructions of the five proteasome assemblies resolved from mature bovine spermatozoa, including PA200-20S (C), free 20S core particle (D), PA200-20S-PAC1/2 (E), PAC1/2-20S-PAC1/2 (F), and PAC1/2-20S (G). Side and a representative top views are shown. (H) Segmented crown-shaped density fitted with PAC1/2 model, showing strong agreement between them and high resolution structrural features. (I) For the three PAC1/2-bound structures, top and bottom views of the engaged 20S, consistently showing absence of the α2 subunit at all PAC1/2-occupied interfaces. (J) Relative abundance of the five proteasome assemblies in mature sperm.

To overcome the low abundance of endogenous sperm proteasomes on cryo-EM grids, we adapted and simplified our IAAG on-grid affinity strategy^51^ by functionalizing graphene oxide-lacey carbon grids with the PYR-NHS linker alone to enhance particle concentration. Single-particle cryo-EM resolved five proteasome configurations at 2.76-3.18 Å resolution (Fig. S2, S3). Two corresponded to the expected PA200-20S and free 20S species (Fig. 1C-D). The other three contained a crown-shaped density, distinct from PA200 and the 19S regulatory particle, capping one or both ends of the 20S or paired with PA200 on the opposite end (Fig. 1E-G). MS and model-to-density fitting identified this crown density as the assembly-chaperone PAC1-PAC2 (PSMG1-PSMG2) heterodimer (Fig. 1A, H), defining three unexpected assemblies: PA200-20S-PAC1/2, PAC1/2-20S-PAC1/2, and PAC1/2-20S (Fig. 1E-G), occupying 41% of the particle population (Fig. 1J). No PAC1/2-bound proteasomes were detected in bovine testis preparations purified by the same strategy (Fig. S4; discussed below), indicating that these species are specific to mature spermatozoa.

These sperm PAC1/2-bound proteasomes differ fundamentally from previously reported PAC1/2-bound assembly intermediates^32,35,36,42^. At every PAC1/2-engaged end, the α-ring lacks the α2 subunit (Fig. 1I), whereas all reported PAC1/2-bound 20S assembly intermediates retain a complete seven-member α-ring^32,35,36,42^. Notably, PAC1/2 also binds in a distict orientation on the CP (Fig. 3A; detailed below) and, unlike transient intermediates that require engineered constructs, cross-linking or specialized capture^32,35,36,42^, the sperm PAC1/2-20S complexes are sufficiently stable for endogenous purification. These features indicate that PAC1/2 has repurposed from an assembly chaperone and may adopt a non-canonical role.

### PAC1/2-bound sperm proteasomes are selectively enriched for α4s

Although α4 and α4s share high sequence identity, ten selective diagnostic residues could distinguish the paralogs in high-reslution cryo-EM density (Fig. S5A). Inspectation of these positions showed that the PAC1/2-20S-PAC1/2 density resolved at 2.88-Å resolution fits α4s model better than α4 at nine of the ten sites (Fig. 2A, D), indicating near-homogeneous α4s incorporation. By contrast, PA200-20S and free 20S maps showed mixed α4s/α4 signatures, with α4s favoured at six positions and α4 or ambiguous density at the remaining sites (Fig. 2B-D).

**Fig. 2.**
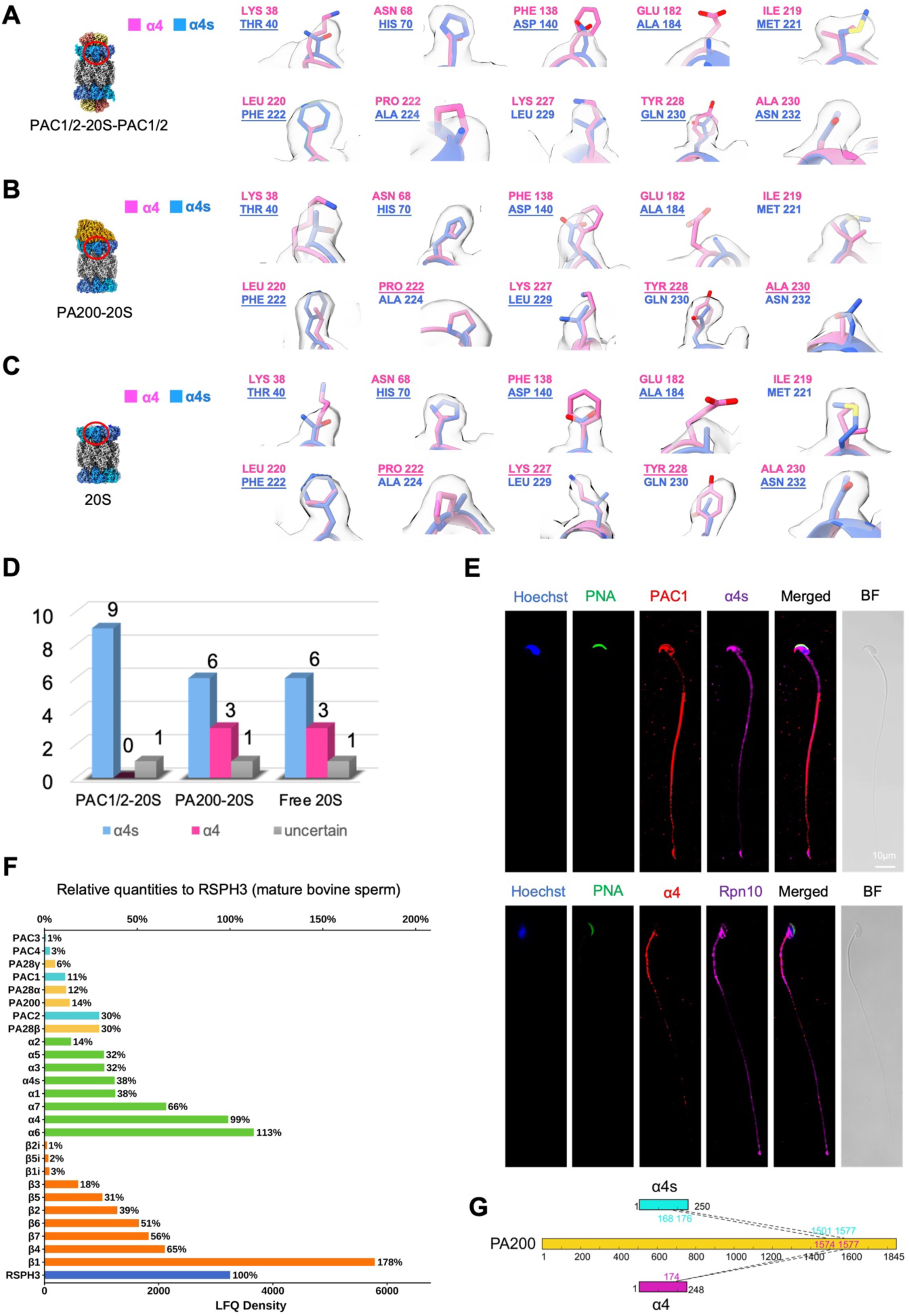
α4s stratifies subunit usage across sperm proteasome assemblies. (A-C) Model-to-density comparisons at ten representative distinct α4/α4s positions in PAC1/2-20S-PAC1/2 (A), PA200-20S (B), and free 20S (C). The α4 model is shown in hot pink and α4s in royal blue; underlined residues donote the better-fitting model for each density. (D) Quantification of α4 and α4s assignments across the three assemblies. (E) Immunofluorescence of mature mouse spermatozoa showing the subcellular localization of PAC1, α4s, α4, and RPN10, a representative 19S subunit. Hoechst (Hoechst33342) labels the nucleus; PNA marks the acrosome; BF, bright field. Scale bar:10μm. (F) Label-free quantitative mass spectrometry (LFQ-MS) of total bovine sperm lysate. The abundance of 20S subunits, assembly chaperones, and ATP-independent activators are normalized to a common radial spoke head component RSPH3 (set to 100%). (G) Cross-linking mass spectrometry (XL-MS) showing direct crosslinks between α4s or α4 and PA200 in sperm proteasome.

Orthogonal analyses supported these structural assignments. Immunofluorescence assay confirmed the presence of both α4s and α4 in mature mouse and bovine spermatozoa and also revealed close PAC1– α4s co-localization in the nuclear compartment and along the sperm flagellum (Fig. 2E, Fig. S1G). By contrast, α4 was predominantly enriched in the mitochondrial midpiece of the flagellum, whereas the 19S subunit Rpn10 (PSMD4) was detected in the post-acrosomal sheath and the flagellum (Fig. 2E, Fig. S1G). Label-free quantitative MS (LFQ-MS) of total bovine sperm lysate^48^ detected both α4s and α4 (Fig. 2F), and cross-linking mass spectrometry (XL-MS) captured distinct α4s-PA200 and α4-PA200 crosslinks in mature sperm (Fig. 2G). Thus, PAC1/2-bound sperm proteasomes are essentially α4s-exclusive, whereas PA200-20S and free 20S are mixed populations. This selective PAC1/2-α4s coupling complements human sperm cryo-ET data, which detected an α4s splice variant with 20S and PA200-20S, but no PAC1/2-bound states, in nuclear lacunae^47^, suggesting that PAC1/2-bound proteasomes may occupy a distinct sperm compartment.

### Sperm PAC1/2-20S complexes are architecturally distinct from canonical assembly intermediates

To determine how sperm PAC1/2-bound proteasomes relate to assembly intermediates, we superimposed the bovine sperm PAC1/2-20S-PAC1/2 structure onto the human PAC1/2-20S-PAC1/2 preholo intermediate (PDB: 8QYS)^32^. In the sperm complex, PAC2 tilts downward by ∼34° (27 Å) toward the vacant α2 position, accompanying by an ∼18 Å lateral translation of PAC1/2 (measured at the PAC1/2 center), which repositions the heterodimer centrally over the α-ring rather than asymmetrically over the α5-α7 hemisphere as in the human intermediate (Fig. 3A)^32,36^. This repositioning is stabilized by two newly ordered interface loops: PAC2 loop 166-177 makes extensive H-bound/salt bridges contacts with α3 through electronstatic and hydrophilic interactions (Fig. 3B-C), and PAC1 loop 107-119 engages α4s and α5 through extensive H-bound/salt bridges network (Fig. 3D). Both loops are disordered in the human assembly intermediate^32^. These mature-state contacts also differ from the yeast Pba1 81–117 loop, which inserts into the α3-α4 pocket in all early precursor complexes (PCs)^35^, reinforcing that sperm PAC1/2 has adopted a non-canonical role.

**Fig. 3.**
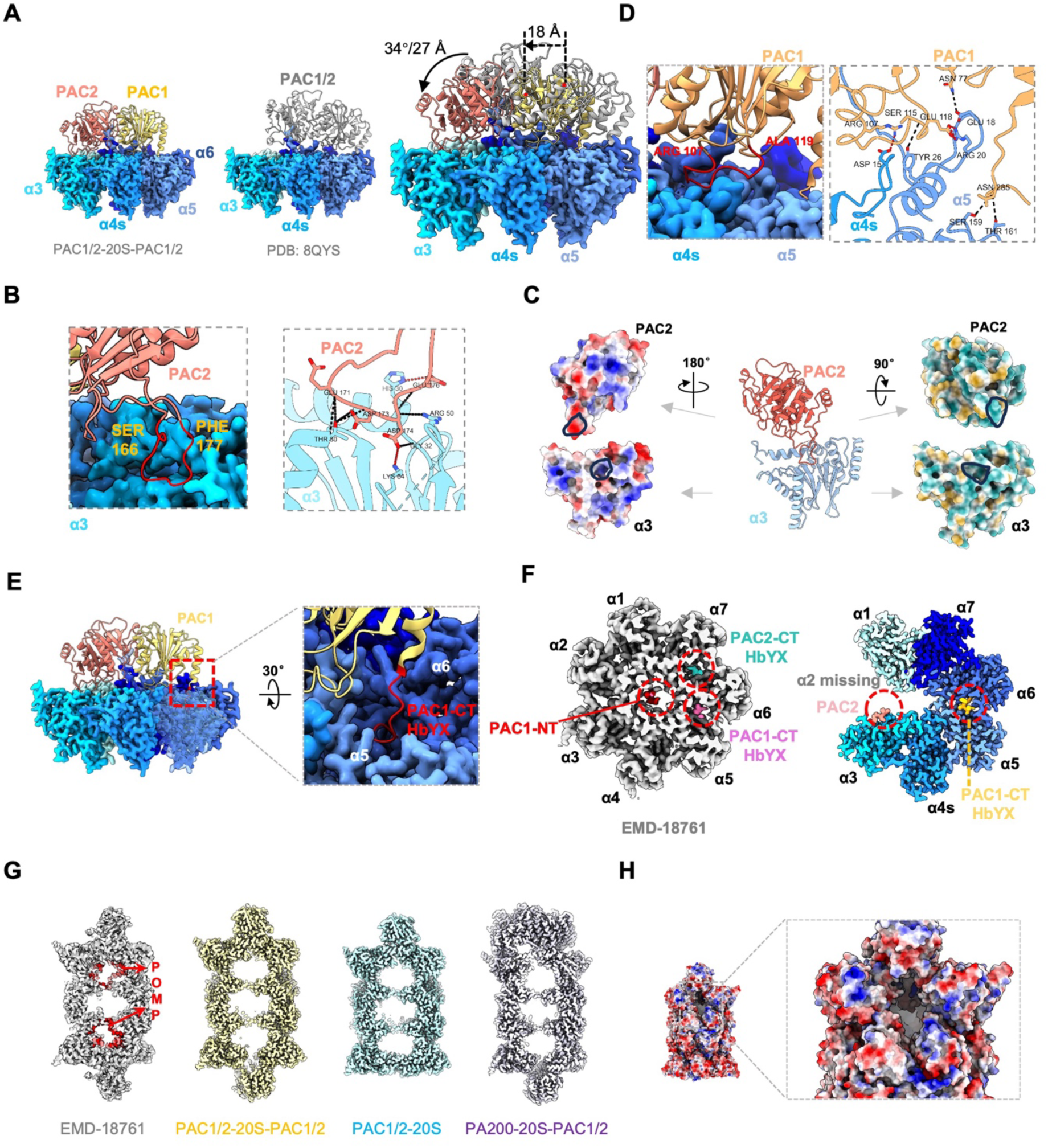
Distinct binding mode, interaction interfaces and POMP-free mature state of mature sperm PAC1/2-20S proteasome assemblies. (A) Structrual Comparison with the human preholo 20S CP (PDB: 8QYS) reveals a downward rotation of PAC1/2—most pronounced for PAC2 (34°/27 Å)—toward the 20S α-ring, together with an ∼18 Å lateral shift toward the vacent α2 position, repositioning PAC1/2 from the α6 side to the center of the 20S. (B) Close-up of the PAC2-á3 interaction interface, shown as a surface representation with a residue-level interaction network. (C) Electrostatic (left) and hydrophobicity (right) surface properties of the PAC2-α3 interface; black outlines indicate reciprocal binding footprints. (D) Close-up of the PAC1-α4s/α5 interface. (E) Rotated view of PAC1/2 docked onto the α-ring, highlighting insertion of the PAC1 C-terminal HbYX motif into the α5/α6 pocket. (F) Comparison of PAC1/2-binding sites in the preholo 20S CP (left) and sperm PAC1/2-20S-PAC1/2 complex (right); binding sites are outlined with red dashed circles. (G) Central slices of the preholo 20S CP (left) with PAC1/2-bound sperm proteasomes (right). POMP (red) is prsent only in the preholo 20S CP chamber, but absent from all PAC1/2-bound sperm proteasome assemblies. (H) Electrostatic surface properties of the PAC1/2-bound proteasome at the α2-deficient position.

HbYX-motif engagement is also remodelled. In human assembly intermediates, the C-terminal HbYX motifs of PAC1 and PAC2 occupy the α5-α6 and α6-α7 pockets, respectively, while the PAC1 N terminus occludes the otherwise open 20S gate (Fig. 3F)^32^. By contrast, in the sperm structure, the PAC1 HbYX motif remains engaged in the α5-α6 pocket (Fig. 3E-F), whereas the PAC2 C-terminal segment (residues 258–264) is disordered and no longer occupies the α6-α7 pocket. The PAC1 N terminus is likewise disordered and no longer blocks the 20S gate (Fig. 3F). Together, these changes leaves the gate open and unobstructed, and thus compatible with substrate entry.

Furthermore, the 20S CP in these complexes is fully mature. In all reported PAC1/2-20S assembly intermediates, the maturation factor POMP occupies the 20S chamber (Fig. 3G), and its autocatalytic degradation by the newly activated β-ring is the canonical event that triggers PAC1/2 release and marks completion of CP maturation^32,36,42–45^. Strikingly, POMP density is absent from all three sperm PAC1/2-bound structures (Fig. 3G), and the β-propeptides, including that of β2, are fully processed (Fig. S5B), establishing that the bound 20S is fully mature rather than an arrested precursor^43^.

### The testis proteasome landscape is dominated by PA200-20S and lacks PAC1/2-bound species

To determine whether the sperm-specific PAC1/2-bound configurations also exist in testicular germ cells, we purified endogenous proteasomes from bovine testis using the same protocol applied to mature sperm. Cryo-EM resolved PA200-20S at 2.97 Å and free 20S at 2.56 Å, but no PAC1/2-associated proteasomes were detected (Fig. 4A, B, Fig. S4). Thus, the α2-deficient, PAC1/2-bound architecture is not a prevalent testicular state but emerges during terminal sperm maturation. In testis PA200-20S, PA200 forms an asymmetric HEAT-repeat solenoid on one CP end and opens the engaged α-ring gate, while the distal gate remains closed (Fig. 4A), consistent with previous PA200-20S structures^12,52,53^ and a recent native human sperm 20S-PA200 structure^47^. Moreover, our structures reveal a gate plasticity for the α4s-containing PA200-20S, specifically in the N-terminal regions of α4s and α3 (discussed in detailed below; Fig. S5C).

**Fig. 4.**
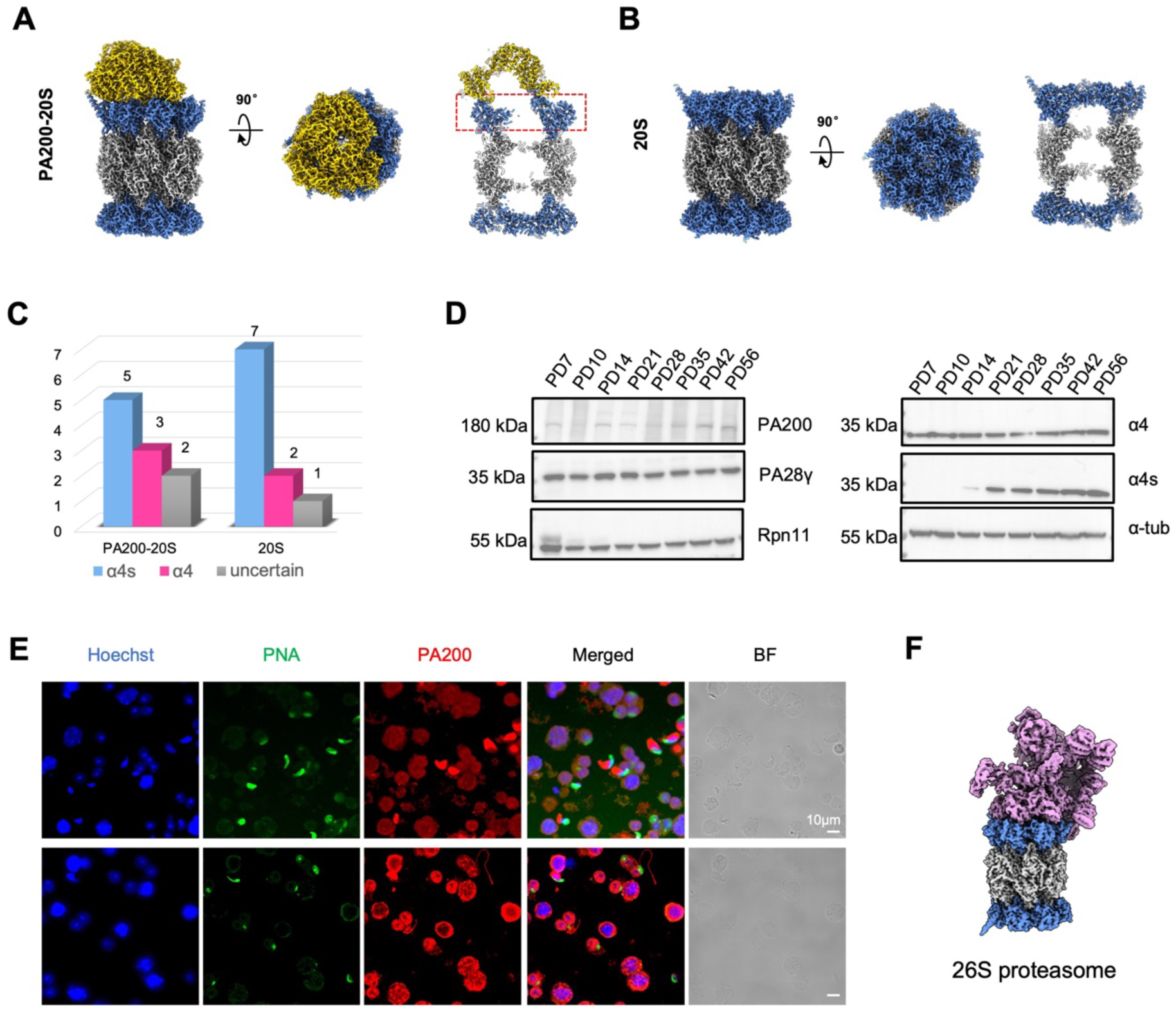
Testis proteasome landscape and charaterization. (A) Cryo-EM structure of the bovine testis PA200-20S proteasome, showing asymmetric PA200 binding to the 20S core and partially opening the engaged 20S gate. (B) Cryo-EM structure of the testis free 20S. (C) Quantification of α4 versus α4s assignment in PA200-20S and free 20S based on the 10 diagnostic residues. (D) Western blot showing the expression patterns of PA200, Rpn11, PA28γ, α4s, and α4 in mouse testis lysate. Tubulin, loading control. (E) Immunofluorescence of PA200 localization in mouse testicular germ cells. Hoechst (Hoechst33342) labels the nucleus; PNA marks the acrosome; BF, bright field. Scale bar:10μm. (F) Cryo-EM structure of the bovine testis 26S proteasome.

Model-to-map fitting at the α4/α4s position revealed a mixed α4s/α4 composition in testis PA200-20S, whereas free 20S preferentially incorporates α4s (Fig. 4C, Fig. S4E-F). To place these structural observations in a developmental context, we raised an in-house monoclonal antibody specific for mouse α4s (with no detectable cross-reactivity to α4) and profiled mouse testis lysates across postnatal development by western blot. α4 was detectable from postnatal day 7, whereas α4s appeared at day 14, coincident with the emergence of pachytene spermatocytes (Fig. 4D). Published single-cell RNA-seq data from staged mouse testicular germ cells confirmed upregulation of *Psma8* (the gene code the subunit α4s) from the pachytene stage^54^ (Fig. S5D), and immunofluorescence showed broad PA200 expression across adult mouse testicular germ cell stages (Fig. 4E). Together, these data support a meiotically timed α4-to-α4s transition^18,25,26^, and place PAC1/2-bound, α2-deficient proteasomes as a post-testicular, terminal-maturation event.

### 26S proteasomes are a minor species in testis and near absent from mature sperm

Across independent mature-sperm purifications, 26S proteasomes were detected only rarely by NS-EM (Fig. S1E-F), insufficient for structural analysis. LFQ-MS nevertheless detected relatively low abundant of 19S subunits in total bovine sperm lysate (Fig. S1H), indicating that 19S components are present in mature sperm but largely fail to assemble into 26S holoenzymes. This is consistent with the reduced ATP pool of cauda epididymal and ejaculated spermatozoa^55^, which would disfavor ATP-dependent 19S loading, and with recent cryo-ET of human sperm, in which 26S particles were not observed in nuclear lacunae^46,47^.

Sufficient 26S proteasome were obtained from testis samples for cryo-EM analysis (Fig. S6A-B). In the final particle pool, 26S proteasomes represented 27.6% of the population, compared with 15.7% for PA200-20S and 56.6% for free 20S (Fig. S6C). The testis 26S proteasome was resolved in the resting state at 3.84 Å, with its 20 CP locally refined to 3.10 Å (Fig. 4F, Fig. S6B). Density at the α4/α4s position favored α4s over α4 (Fig. S6D-E). Together, the near absence of 26S proteasomes in mature sperm indicates that ATP-independent proteolysis—mediated by PA200-20S, and potentially also by PAC1/2-20S and PA200-20S-PAC1/2—is the principal proteolytic pathway in mature sperm. This contrast sharply with somatic cells, in which the 26S proteasome predominates, whereas in testis both ATP-dependent and ATP-indepent assemblies appear to contribute to proteolysis.

### α4s-containing proteasomes are assembled through the canonical chaperone pathway and α2 exclusion is dictated by the mature-sperm environment

The coexistence of α4s– and α4-containing proteasomes in testicular germ cells and mature sperm raised two questions: wheter α4s uses the canonical PAC1-PAC4/POMP assembly chaperones and follow the same pathway, and whether α4s reshapes the intrinsic catalytic perperties of the 20S sCP. To address both, we replaced endogenous α4 in HEK293F cells with stably expressed human HA-α4s, and analysed HA-immunoprecipitates by MS (Fig. 5A-B, Fig. S7A). The HA-α4s interactome matched the HA-α4 controls, recovering all 20S subunits, PAC1-PAC4, POMP, PA200, and 19S components (Fig. 5B). Western blotting confirmed co-association of PAC1, PAC2, POMP, PA200, and the 19S subunit RPN6 with α4s-containing particles in both human and mouse settings (Fig. 5C, Fig. S7B).

**Fig. 5.**
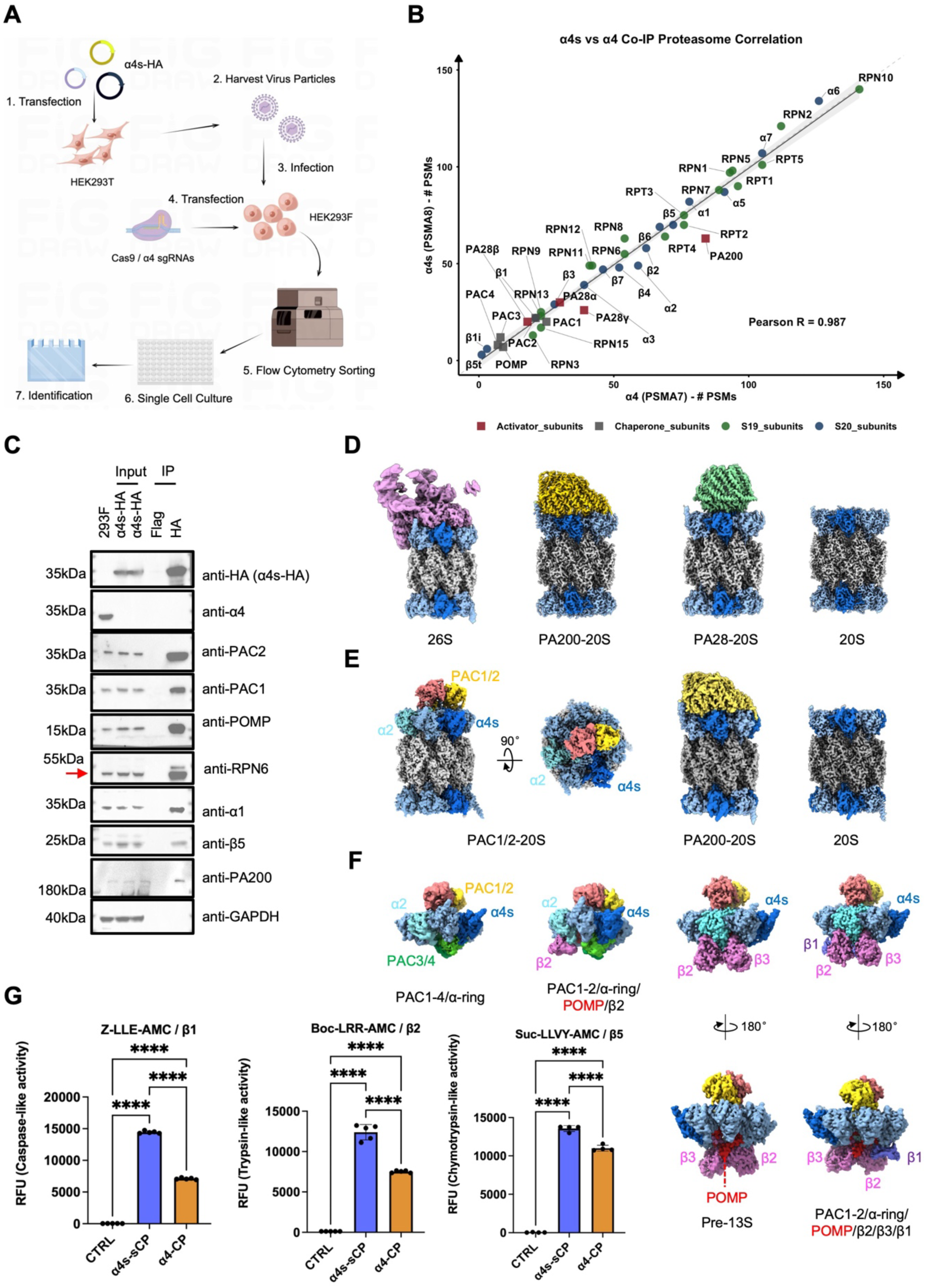
α4s-containing proteasomes assemble through the canonical chaperone pathway yet are intrinsically hyperactive. (A) Schematic of the strategy for generating α4-knockout HEK293F cells stably expressing α4s-HA. (B) Correlation of HA co-immunoprecipitation-mass spectrometry datasets from murine α4s-HA (α4 knockout) versus murine α4-HA cells. (C) Western blot validation of proteasome subunits and canonical assembly chaperones, including α4, α1, β5, RPN6, PA200, PAC1, PAC2, and POMP. GADPH, loading control. (D) Cryo-EM structures of human α4s-containing proteasome assemblies purified from α4 knockout HEK293F cells overexpressing HA-tagged α4s, including 26S proteasome with a highly dynamic 19S lid, PA200-20S, PA28-20S, and mature 20S CP. (E-F) Cryo-EM structures of human α4s-containing proteasomes (E) and α4s-containing assembly intermediates (F), purified from α4 knockout HEK293F cells expressing HA-tagged α4s and further co-expressing His-tagged PAC1 and Flag-tagged PAC2 via tandem HA and Flag affinity purification. Both analyses show that the reconstituted PAC1/2-bound α4s-containing sCPs in HEK293F cells retained α2. (G) Chymotrypsin-like, trypsin-like, and caspase-like peptidase activities of purified mature human α4s-sCP or α4-CP, measured with Suc-LLVY-AMC, Z-LLE-AMC, and Boc-LRR-AMC, respectively. Data are mean ± s.d. of n = 4 or 5 technical repeats; statistical analysis was performed using one-way ANOVA with Tukey’s multiple-comparisons test; \*\*\**p* < 0.001, and \*\*\*\**p* < 0.0001.

Cryo-EM of the HA-purified α4s-containg complexes resolved the mature 20S sCPs, the 26S proteasome with an extremely dynamic lid, PA200-20S, and PA28-20S at the 2.47-3.07 Å resolution range, together with three discrete assembly intermediates (Fig. 5D, Fig. S8A). The intermediate maps showed PAC1-PAC4 and POMP occupying the same positions on nascent α4s-containing α-rings as on constitutive α4-containing intermediates^32,36,42^ (Fig. S8A). A similar set of assembly intermediates was obtained from α4-knockout HEK293F cells expressing mouse α4s (Fig. S9A). In direct contrast to mature spermatozoa, no PAC1/2-bound mature 20S complexes were detected in either cellular context. Collectively, these data establish that α4s is incorporated into mature 20S CPs through the canonical PAC1-PAC4/POMP chaperone systems.

The absence of the α2 from every mature-sperm PAC1/2-engaged α-ring raised a further question: does α2 exclusion arise intrinsiclly from PAC1/2 binding to α4s-containing 20S sCPs, or is it instead imposed by the mature-sperm cellular context? To address this, we replaced endogenous α4 in HEK293F cells with stably expressed α4s-HA and co-expressed His-PAC1 and Flag-PAC2 (Fig. S10A). Sequential anti-HA and anti-Flag affinity purification yielded PAC1/2-bound 20S sCPs, PA200-sCP, and free 20S sCP (Fig. 5E), together with a series assembly intermediates (PAC1-4/α-ring, PAC1-2/α-ring/POMP/β2, Pre-13S, and PAC1-2/α-ring/POMP/β2/β3/β1), all of which were resolved by cryo-EM (Fig. 5F, Fig. S10B-C). Notably, the PAC1-2/α-ring/POMP/β2/β3/β1 complex represents a previously uncharacterized assembly intermediate, suggesting a potentially distinct assembly route with β2-β3-β1 incorporation ordering, rather than the cononical β2-β3-β4 sequence, while still sharing the same chaperone system. Unlike the sperm complexes, however, these reconstituted particles retained α2 (Fig. 5E-F). Thus, α2 exclusion is driven neither by α4s incorporation nor by PAC1/2 engagement alone, but instead requires the mature-sperm cellular environment, implicating sperm-specific post-translational modifications, disassembly factors or proteolytic remodelling during late sperm maturation.

### α4s-containing proteasomes are intrinsically hyperactive

Closer inspection revealed that α4s incorporation into the sCP remodels the electrostatic surface relative to α4 in canonical CP (Fig. S11A-B). In the top view, α4s contains an N-terminal Arg in place of the corresponding Ser in α4, creating a stronger positive potential at the subsurface α-ring gate region (Fig. S11A). In addition, the α4s loop that engages α3 (residues 49-57) is more positively charged than the corresponding α4 loop (residues 47-55) (Fig. S11A). In the side view. In addition, an α3-adjacent flank region in α4s (residues 216-232) displays a more positive or neutral electrostatic character, whereas the corresponding region in α4 is more negative, consistent with sequence divergence between α4s and α4 in this region (Fig. S11B).

To investigate whether these structural differences translate into altered catalytic capacity, we purified free 20S from HA-α4s and HA-α4 HEK293F cell lines (Fig. S7C-E), and measured chymotrypsin-like (β5), caspase-like (β1), and trypsin-like (β2) activities using fluorogenic substrates. Human α4s-containing sCP exhibited higher basal activity at all three catalytic sites than α4-containing controls, with increases of 1.23-, 2.03-, and 1.65-fold for β5, β1, and β2, respectively (Fig. 5G). This enhancement was conserved in mouse α4s-containging 20S (Fig. S7F). Thus, incorporation of α4s consistently increases all three peptidase activities, with a more pronounced effect at β1 in both species, suggesting a particular specialization for acidic-residue substrates. This broad increase in proteolytic activity is likely to reflect an intrinsic, structure-encoded property of the mature α4s spermatoproteasome. Together, these data establish the spermatoproteasome as a purpose-built, hyperactive proteasome variant adapted to the elevated proteolytic demands of sperm differentiation, maturation, and fertilization.

## Discussion

Our cryo-EM analysis of endogenous mature bovine sperm proteasomes reveals an unexpectedly diverse, ATP-independent proteasome landscape. Beyond PA200-20S and free 20S, we identify three mature-sperm-specific assemblies—PAC1/2-20S, PAC1/2-20S-PAC1/2, and a previously unrecognized PA200-20S-PAC1/2 hybrid—that are mature, POMP-free, α4s-enriched, and α2-deficient. Their existence extends the prevailing view, supported by all assembly intermediate characterized to date^32,33,35–37,42–45^, that PAC1/2 is solely a transient assembly chaperone released upon CP maturation. Instead, mature sperm repurpose PAC1/2 as a stable post-assembly regulator of a specialized 20S proteasome state in terminally differentiated male gametes. Reconstitution in HEK293F cells further shows that α2 exclusion is imposed by the mature-sperm environment rather than by α4s or PAC1/2 alone, whereas α4s-containing proteasomes are assembled through the canocial PAC1-PAC4/POMP chaperone system yet may follow the β2-β3-β1 incorporation ordering rather than the known β2-β3-β4 ordering.

Several structural features distinguish the sperm PAC1/2-bound state from an assembly intermediate. First, all PAC1/2-bound sperm complexes lack POMP and contain fully processed β-propeptides (Fig. 3G, Fig. S5B), excluding an immature precursor identity^35,42–45^. Second, PAC1/2 is repositioned by a ∼34° tilt and an ∼18 Å shift toward the α2 vacancy; the PAC2 C terminus is released from the α6-α7 pocket, while new PAC2-α3 and PAC1-α4s/α5 interfaces are stabilized (Fig. 3A, B, D). Third, the PAC1 N terminus is disordered rather than gate-occluding^32,36^, leaving the gate open and unobstructed (Fig. 3F). Consistent with this architecture, and combined with the widened substrate entry channel caused by the α2 vacancy, bovine sperm proteasomes display markedly elevated activity across all three catalytic specificities relative to testis or muscle proteasomes (Fig. S11C). We therefore propose that, in mature sperm, PAC1/2 functions as an ATP-independent, gate-opening activator that sustain proteolytic capacity after biosynthesis has largely ceased. The PA200-20S-PAC1/2 hybrid extends this principle by placing two ATP-independent regulators with distinct substrate preferences on opposite ends of one sCP, potentially coupling PA200-dependent processing of acetylated nuclear proteins with broader PAC1/2-stimulated proteolysis.

How mature-sperm selectively remove α2 remains an important mechanistic question. One plausible route involves the PI31-Fbxo7 axis: PI31 is strongly enriched with sCPs together with its stablizing partner Fbxo7^22^, both of which are essentail for spermatogenesis^56,57^. Because PI31 contacts multiple α-subunits but not α2 or α5^58^, and Fbxo7 can ubiquitinate α2^59^, this axis could weaken or mark α2 for displacement from the α4s-containing α-ring. Alternatively, sperm-specific post-translational modifications (PTM) may exploit the intrinsic fragility of the α1-α2 junction—the only α-α interface lacking the conserved Y-D hydrogen bond^60^, with α2 carring the densest PTM landscape among α-subunits^61^— to destabilize α2 anchorage to α1/α3^62^. The retention of α2 in reconstituted α4s-containing sCPs indicates that α2 eviction requires a mature-sperm-specific second inut, positioning PAC1/2-bound assemblies as post-maturation proteasomes poised for rapid activation upon capacitation.

Our structures also reveal a compositional stratification of the sperm proteasome pool: PA200-20S and free 20S are mixed α4s/α4 assemblies in which α4s is more abundant, mirroring their testicular counterparts, whereas PAC1/2-bound complexes are essentially α4s-exclusive (Fig. 2D). During spermatogenesis, α4 is detected from postnatal day 7 and α4s at day 14 (Fig. 4D), a pattern mirrored at the transcript level^54^, supporting a developmental shift toward α4s-containing proteasomes. α4s-specific co-immunoprecipitation followed by LFQ-MS recovered abundant α4s-unique peptides without obvious enrichment of α4-unique peptides (Fig. S11D), in agreement with previous biochemical observations^24^, and disfavor α4s/α4-hybrid CPs containing different paralogs on opposite α-rings. Thus, α4 and α4s appear to form biochemically segregated, rather than mosaic, 20S populations. Furthermore, our structures reveal a gate plasticity for the α4s-containing PA200-20S. Within α4s-containing PA200-20S, the activated α-ring face is also asymmetrically dynamic: in bovine testis and mature-sperm PA200-20S maps, and in the human PA200-sCP map, α3 loses density throughout its N-terminal H0 helix, reverse turn and YDX motif, whereas α4s retains only a partial H0 helix and entirely lacks density for the extended reverse turn, YDX motif and 120-124 loop (Fig. S5C). This α4s-α3 gate plasticity may help tune substrate entry in the rapidly changing proteolytic enviroment of post-meiotic sperm.

Despite this specialization, α4s-containing 20S sCPs are assembled by the canonical PAC1-PAC4/POMP chaperone system (Fig. 5B-C, F), yet are intrinsically hyperactive at all three catalytic sites (∼1.2–2-fold in human; Fig. 5G, Fig. S7F). Hyperactivity is therefore a structure-encoded property of the mature spermatoproteasome, not a consequence of an alternative assembly route. Our structures also capture a previously uncharacterized PAC1-2/α-ring/POMP/β2/β3/β1 intermediate (Fig. 5F), providing mammalian structural evidence for a β2→β3→β1 incorporation order previously inferred from a biochemical study^63^. Whether this ordering is α4s-coupled or reflects a broader flexibility in mammalian 20S biogenesis remains to be explored.

Mechanistically, α4s remodels the electrostatic landscape of the the α-ring, including the ourter surface, inter-subunit loop and inner pore, with enhanced positive electrostatic potential near the active centers of β1 and β5 (Fig. S7G). Such remodelling is predicted to favour engagement of acidic, phosphorylated and hydrophilic substrates that accumulate during spermiogenesis and capacitation, consistent with preferential enhancement of β1 activity, which points to a specialization for acidic-residue substrates. The α4-to-α4s switch, coincident with pachytene entry around postnatal day 14^22,54^ (Fig. 4F, Fig. S5B), therefore represents a meiotically gated commitment that installs a hyperactive, substrate-tuned proteasome in the post-meiotic germline. Defineing the factors that license α4s selection by PAC1-PAC4/POMP will be central to understanding how the spermatoproteasomes are configured during mammalian gametogenesis.

Taken together, these findings place ATP-independent proteolysis at the centre of the male-germline proteasome landscape. Although 26S proteasomes remain moderate in testis, they are nearly absent from mature spermatozoa despite the persistence of 19S subunits in lysate, consistent with the low ATP pool of mature sperm^55^, which would disfavour 26S assembly. Our bovine structures broadly agree with recent cryo-ET of human spermatozoa^47^, which identified 20S and PA200-20S as dominant nuclear species in lacunae, but did not detect PAC1/2-bound states, suggesting that these complexes may be compartmentally distinct, species-specific or maturation-state-dependent. Spatially resolved mapping will be needed to define where each proteasome variant acts and which substrates it processes during capacitation, acrosomal exocytosis and zona pellucida penetration^21,50,64^.

In summary, we define a sperm-specific, ATP-independent proteasome landscape that expands the functional repertoire of both α4s and PAC1/2. PAC1/2 can act not only as a transient chaperone, but also as a stable post-assembly regulator of a specialized, α2-deficient 20S CP, while α2 exclusion reveals a mature-sperm-specific layer of proteasome regulation. In addition, α4s-containing proteasomes are assembled through the canonical PAC1-PAC4/POMP chaperone system yet may follow distinct β subunit incorporation ordering; α4s generates intrinsically hyperactive proteasome whose α-ring remodelling and altered active center proterty may tune substrate selection during sperm maturation and fertilization. These findings underscore the remarkable plasticity of proteasome regulation in specialized cell types, provide a structural framework for how proteasome specialization supports male gamete function and fertility, and open new avenues for investigating idiopathic infertility and for therapeutic targeting of sperm proteasome specialization.

## Methods

### Mouse

All animal experiments in this study were approved by the Institutional Animal Care and Use Committee of the Center for Excellence in Molecular Cell Science, Institute of Biochemistry and Cell Biology, Chinese Academy of Sciences, and complied with strict ethical guidelines. Male wild-type C57BL/6J mice at 8-10 weeks were purchased from Shanghai SLAC Laboratory Animal Co., Ltd and usded for testicular germ cells and mature sperm isolation. Animals were housed in a specific-pathogen-free (SPF) facility with a 12/12-h light/dark cycle. The temperature was maintained at 22–24 °C, and relative humidity was kept within 45–65%. Husbandry and handling of mice followed a previous protocol^65^.

### Cell lines generation

HEK293F and Flag-PA200-expressing HEK293F cells were transduced with lentivirus encoding HA-tagged human or mouse α4s (α4s-HA) or α4 (α4-HA). Lentiviruses were produced in HEK293T cells by co-transfection of pCDH-CMV-α4s/α4-HA-EF1-coGFP, psPAX2, and pMD2.G using Lipo8000 (Beyotime). Viral supernatants collected at 48 h and 72 h were filtered through a 0.22-µm membrane, and applied to target cells with 8 µg/mL polybrene. Endogenous α4 was disrupted by CRISPR/Cas9 using pX330-sgRNA-mCherry plasmids targeting the first exon of the α4 genome. Double –positive (coGFP^+^ and mCherry^+^) single cells were isolated by FACS, and expression of HA-tagged α4s or α4 and loss of endogenous α4 were verified by immunoblotting.

### Co-immunoprecipitation (Co-IP)

HEK293F cell pellets were resuspended in ice –cold Co-IP lysis buffer (50 mM Tris-HCl pH 7.5, 150 mM NaCl, 10 mM MgCl_2_, 2 mM ATP, 10% Glycerol, 1mM β-ME, 1×PIC) and lysed on ice for 1 h. Lysates were clarified by concentration at 12,000×g for 30 min at 4°C, and the supernatants were incubated overnight at 4°C with 50 μL of epitope affinity matrix. Beads were washed six times with PBST (0.02% Tween 20) containing 2 mM ATP and bound proteins were eluted twice with 500 μL of 0.5 mg/mL HA peptide. Elutes were pooled, concentrated and stored at –80°C until used for western blotting or mass spectrometry analysis.

### Immunofluorescence assay

Immunofluorescence analysis of spermatozoa and testicular germ cells was performed essentially as described^66^. Cells were fixed for 15 min at room temperature with 4% Tissue Fix Solution (T17024; Saint-Bio Biotech), deposited onto coated slides, and permeabilized for 10 min in enhanced immunostaining permeabilization buffer (M20254; Saint-Bio Biotech). Samples were blocked for 1 h in 5% BSA in PBST (0.05% Tween-20) and incubated overnight at 4°C with primary antibodies diluted in 1% BSA solution. After three PBST washes, slides were incubated for 1h at room temperature with AF647-labeled goat anti-mouse IgG (H+L) (A0473; Beyotime) or Goat anti-Rabbit IgG (H+L) Cross-Adsorbed Secondary Antibody, Alexa Fluor™ 555 (A-21428; Thermo Fisher Scientific), each diluted 1:800 in 1% BSA. Peanut agglutinin (PNA; L32458; Thermo Fisher Scientific) was used at a 1:50 in PBST (final concentration, 20 μg/mL). Slides were mounted in antifade mounting medium containing Hoechst 33342 (P0133; Beyotime) and imaged on a Zeiss LSM710 confocal microscope. Antibodies and reagents are listed in the Supplementary Table 4.

### Western blotting

Mouse testes were homogenized in ice-cold lysis buffer supplemented with cOmplete EDTA-free protease inhibitor cocktail (EASYpack 04693132001; Roche Diagnostics GmbH). Lysates were clarified by centrifugation at 12,000×g, mixed with 5 × SDS-PAGE loading buffer, and denatured at 100°C for 10 min. Equal amount of total protein were resolved on FuturePAGE^TM^ 4-20% 15 Wells (ET15420LGel; ACE Biotechnology), transferred to PVDF membranes, blocked with 5% dry nonfat milk in PBST (0.05% Tween 20) at room temperature, and probed overnight at 4°C with primary antibodies followed by secondary antibodies at room temperature. Signals were detected with Western BLoT Ultra Sensitive HRP Substrate (T7104A; Takara) and imaged on ChemiDoc™ MP Imaging System (Bio-Rad). Antibodies and reagents are listed in the Supplementary Table 1.

### Purification of PAC1/PAC2-bound proteasomes from α4-knockout HEK293F cells expressing HA tagged human α4s

α4-knockout HEK293F cells expressing HA-tagged human α4s (α4s-HA) were transiently transfected with pcDNA3.4-PAC1^His^/PAC2^Flag^ plasmids. After 48 h of expression, cells were harvested and lysed by gentle homogenization in ice-cold lysis buffer (50 mM Tris-HCl pH 7.5, 150 mM NaCl, 2 mM ATP, 0.1% NP-40, 10% glycerol, 1×PIC, 1 mM PMSF and 1 mM β-ME) by gentle homogenization, followed by clarification via centrifugation at 12,000 × g for 30 min at 4°C. The soluble supernatant was incubated with pre-equilibrated anti-HA affinity beads (SA068100; Smart-Lifesciences) for 2 h at 4°C with gentle rotation. Beads were washed extensively with wash buffer (50 mM Tris-HCl pH 7.5, 150 mM NaCl, 0.1% NP-40, 10% glycerol, 2 mM ATP,1 mM β-ME), and bound complexes were eluted with wash buffer supplemented with 0.5 mg/mL HA peptide. The HA eluate was incubated overnight at 4°C with anti-DYKDDDDK affinity beads (SA042100; Smart-Lifesciences) overnight at 4°C. After extensive washing with wash buffer (50 mM Tris-HCl pH 7.5, 150 mM NaCl, 1 mM ATP, 0.1% NP-40, 10% glycerol, 1mM β-ME), the PAC1/PAC2-bound proteasome complexes were eluted in PBS (pH7.4) containing 5% glycerol and 0.5 mg/mL Flag peptide. Eluates were concentrated using a 10-kDa-cutoff centrifugal filter at 4°C, aliquoted and stored at –80°C until used for cryo-EM grid preparation. Reagents are listed in the Supplementary Table 4.

### Purification of 20S Proteasomes from α4-knockout HEK293F cells expressing HA-tagged human α4s and Flag-tagged PA200

α4-knockout HEK293F cells expressing HA-tagged human α4s (α4s-HA) and Flag-tagged PA200 were harvested and lysed by gentle homogenization in ice-cold lysis buffer (50 mM Tris-HCl pH 7.5, 150 mM NaCl, 2 mM ATP, 0.1% NP-40, 10% glycerol, 1×PIC, 1 mM PMSF and 1 mM β-ME) by gentle homogenization, followed by clarification via centrifugation at 12,000 × g for 30 min at 4°C. The soluble supernatant was incubated with pre-equilibrated anti-HA affinity beads (SA068100, Smart-Lifesciences, Changzhou) for 2 h at 4°C with gentle rotation. Beads were washed extensively with wash buffer (50 mM Tris-HCl pH 7.5, 150 mM NaCl, 0.1% NP-40, 10% glycerol, 2 mM ATP, 1 mM β-ME), and bound complexes were eluted with wash buffer containing 0.5 mg/mL HA peptide. The HA eluate was further incubated overnight at 4°C with anti-DYKDDDDK affinity beads (SA042100; Smart-Lifesciences). The flowthrough of anti-DYKDDDDK affinity beads was concentrated and loaded onto a Superose 6 Increase 10/300GL column. Eluted fractions were concentrated using a 100-kDa-cutoff centrifugal filter at 4°C, aliquoted and stored at –80°C until used for NS-EM and cryo-EM grid preparation. Reagents are listed in the Supplementary Table 4.

### Proteasome activity assay

Proteasomes or lysates were incubated at 37 °C for 30 min in 50 µL reaction buffer (50 mM Tris-HCl pH 7.5, 150mM NaCl, and 10mM MgCl_2_). Chymotrypsin-like, trypsin-like, and caspase-like activity were measured using Suc-LLVY-AMC, Boc-LRR-AMC, and Z-LLE-AMC, respectively, at a final substrate concentration of 50 μM or 100 μM. AMC release was monitored at 380 nm excitetation and 460 nm emmision) was monitored on a Synergy Neo plate reader(Biotek).

### Statistical analyses

Statistical analysis for proteasome activity assays were conducted using GraphPad Prism software 10.6.1 (GraphPad Software). For each fluorescent substrate, comparisons among the three experiment groups were performed by one-way analysis of variance (ANOVA), followed by Tukey’s multiple comparisons test to detect significant pairwise differences. Data are presented as mean ± standard deviation (SD) with overlaid individual data points overlaid. *P* < 0.05 was considered as statistical significant, siginificance leveles are indicated as:***, *P <0.001*; ****, *P <0.0001*.

### Isolation of mature bovine spermatozoa

Cauda epididymides were dissected from bovine testes and incised to release spermatozoa into PBS solution. The suspension was cleared by sequential centrifugation at 100×g for 1 min and 500×g for 10 min, followed by an additional 500×g wash for 10 min. The pellet was resuspended in PBS, centrifuged at 3000×g for 10 min, and mixed 1:1:1 with ART-2040 and ART-2080 density-gradient media. After centrifugation at 3000×g for 10 min, the lower sperm-enriched fraction was collected, separated from connective tissue and interface material, and stored at –80 °C.

### PA200-20S and 20S proteasome purification from mature bovine spermatozoa and testis

Mature bovine spermatozoa were disrupted using a freezing mixer ball mill and lysed for 5h in 50 mM HEPES pH 8.0, 100 mM NaCl, 5 mM MgCl_2_, 10% glycerol, 0.5% Triton X-100, 1 mM ATP, 1 mM DTT, and 0.1% Supernuclease S. Lysates were clarified by centrifugation at 20,000×g for 1h at 4 °C, and the supernatant was filtered through a 0.45-μm membrane. The filtrate was loaded onto a Source^TM^ 15Q ion exchange column pre-equilibrated with buffer A (50 mM HEPES pH 8.0, 5 mM MgCl_2_, 5% glycerol, 1 mM ATP, 1 mM DTT) contating 200 mM NaCl. The Source 15Q column was washed with buffer A with 200 mM NaCl and eluted with a 400 mL gradient of 200-480 mM NaCl in buffer A. Fractions were assayed for proteasome distribution and activity using peptidase activity assay (described below). Peptidase-active fractions were pooled, concentrated, and further purified via glycerol-gradient-centrifugation (10-40% glycerol (w/v) in 50 mM HEPES pH8.0, 20 mM NaCl, 10 mM MgCl_2_, 1 mM ATP, 1 mM DTT) by centrifugation at 37,000 rpm for 16 h. Proteasome-containing fractions were identified by peptidase assay, SDS-PAGE, and negative-stain EM. Purified samples were flash-frozen in liquid nitrogen and stored at –80°C.

Bovine testes were processed using the same protocol as described for mature spermatozoa. Testis was homogenized and lysed for 5 h in the same lysis buffer (50 mM HEPES pH 8.0, 100 mM NaCl, 5 mM MgCl₂, 10% Glycerol, 0.5% Triton X-100, 1 mM ATP, 1 mM DTT, and 0.1% Supernuclease S). Lysates were clarified by centrifugation at 20,000×g for 1 h at 4 °C, filtered through a 0.45-μm membrane, and applied to a Source 15Q anion-exchange column pre-equilibrated with buffer A containing 200 mM NaCl. Elution, fraction analysis, concentration, and glycerol gradient centrifugation were performed as described above. Final purified samples were flash-frozen in liquid nitrogen and stored at –80 °C.

### Purification of 26S proteasomes from mature bovine spermatozoa and testes

Mature bovine cauda epididymal spermatozoa were disrupted using a freezing mixer ball mill and lysed for 9 h in the lysis buffer (50 mM HEPES pH 8.0, 100 mM NaCl, 5 mM MgCl₂, 10% glycerol, 1% Triton X-100, 2 mM ATP, 1 mM DTT, and 0.1% Supernuclease S). Lysates were clarified by centrifugation at 20,000×g for 1 h at 4 °C, and then filtered through a 0.45-μm filter. Unlike the testes protocol, the filtrate was loaded directly onto a Source 15Q anion-exchange column pre-equilibrated with 20% buffer B (50 mM HEPES pH 8.0, 1M NaCl, 5 mM MgCl₂, 5% glycerol, 1 mM ATP, and 1 mM DTT) supplemented with 80% buffer A (50 mM HEPES pH 8.0, 5 mM MgCl₂, 5% glycerol, 1 mM ATP, and 1 mM DTT). The column was washed with mixed buffer containing 200 mM NaCl and eluted with a 400 mL gradient of 200-480 mM NaCl in mix buffer. Proteasome-containing fractions were pooled, concentrated, and subjected to glycerol gradient centrifugation (10-40% glycerol (w/v), 50 mM HEPES pH 8.0, 50 mM NaCl, 10 mM MgCl₂, 1 mM ATP, 1 mM DTT) at 37,000 rpm for 6 h. Purified 26S proteasomes were confirmed by peptidase activity assay, SDS-PAGE, and negative-stain EM, flash-freezing in liquid nitrogen and stored at –80 °C.

Bovine testes were peeled to remove the outer tunica, diced and crushed using a freezing mixer ball mill, and lysed for 30 min in lysis buffer (50 mM HEPES pH 8.0, 50 mM NaCl, 10 mM MgCl₂, 10% glycerol, 1 mM ATP, 1 mM DTT). Homogenates were clarified by centrifugation at 20,000×g for 40 min at 4°C, followed by ultracentrifugation at 100,000×g for 1 h. The supernatant was filtered through four layers of gauze and loaded onto a DEAE ion exchange column equilibrated in buffer A (50 mM HEPES pH 8.0, 10 mM MgCl₂, 5% Glycerol, 1 mM ATP, 1 mM DTT). The DEAE column was sequentially washed with buffer A containing 50 mM NaCl and 150 mM NaCl, and then eluted with 100 mL of buffer A containing 150 mM NaCl. The eluate was further purified on a Source 15Q column equilibrated in buffer A containing 150 mM NaCl, washed with the same buffer and eluted with a 500-mL gradient of 150-500 mM NaCl in buffer A. Proteasome-containing fractions were identified by a peptidase activity assay, pooled, concentrated, and further purified by glycerol gradient centrifugation (10-40% glycerol (wt/vol), 50 mM HEPES pH8.0, 50 mM NaCl, 10 mM MgCl_2_, 1 mM ATP, 1 mM DTT) at 37,000 rpm for 6 h. Final proteasome fractions were verified by peptidase activity assay, SDS-PAGE, and negative-stain EM. Samples were flash-frozen in liquid nitrogen and stored at –80°C.

### LC/tandem MS (MS/MS)

Protein samples were precipitated with ice-cold acetone, and pellets waere dried in a SpeedVac for 1-2 min. Pellets were resuspended in 8 M urea containing 100 mM Tris-HCl pH 8.5, reduced with 5 mM TCEP (Thermo Scientific), alkylated with 10 mM iodoacetamide (Sigma), and incubated for 30 min at room temperature. Samples were diluted 4-fold and digested overnight with trypsin (Promega) at a 1:50 enzyme-to-protein ratio (w/w). Digested peptides were desalted on MonoSpin™ C18 columns (GL Science), dried in a SpeedVac and subjected to LC-MS/MS analysis.

Peptides were separated on a custom 30-cm pulled-tip analytical column (75 μm inner diameter) packed with 1.9 μm ReproSil-Pur C18-AQ resin (Dr. Maisch GmbH, Germany), and coupled online to an Easy-nLC 1200 nano-HPLC system (Thermo Scientific). The column was maintained at 55 °C during data acquisition. Peptide separation was performed using a binary mobile phase: buffer A (0.1% formic acid in ultrapure water) and buffer B (0.1% formic acid in 80% acetonitrile). The elution gradient was set as follows: 5-15% buffer B over 0-30 min, 15-25% over 30-75 min, 25-40% over 75-105 min, 40-100% over 105-110 min, and a final 100% buffer B hold for 10 min. The flow rate was kept at 300 nL/min throughout the run.

Data-dependent acquisition (DDA) tandem mass spectrometry was performed on a Q Exactive Orbitrap mass spectrometer (Thermo Scientific). Eluted peptides were ionized by electrospray ionization with a distal spray voltage of 2.2 kV. Each acquisition cycle consisted of one full MS scan (m/z 300-1800), followed by MS/MS fragmentation of the top 20 most intense precursor ions at a normalized collision energy of 28%. Full-scan MS was acquired at a resolution of 70,000 with an AGC target of 3×10⁶. MS/MS scans were collected at a resolution of 17,500, with an isolation window of 1.8 m/z and an AGC target of 1×10⁵. Single microscans were used for MS and MS/MS, with maximum injection times of 50 ms and 100 ms, respectively. Dynamic exclusion was enabled with the following parameters: exclusion of charge states 1, 2, and >8; isotope exclusion activated; and exclusion durations of 5, 10, and 15 s. All LC and MS instrumental operations were controlled by the Xcalibur software (Thermo Scientific).

For cross-linked samples, cross-linked peptide pairs were identified using pLink2 (pFind Team, Beijing) according to a previously described workflow. Database search parameters were defined as follows: trypsin as the digestive enzyme, up to three missed cleavage sites, and 20-ppm mass tolerance for both precursor and fragment ions. Carbamidomethylation of cysteine was set as a fixed modification, and methionine oxidation as a variable modification. Final results were filtered at a spectral-level false discovery rate (FDR) of 5%.

### Label-free quantification mass spectrometry (LFQ-MS)

Protein samples underwent two-step tryptic digestion with trypsin at 37 °C, and the reaction was terminated using 20% TFA. We then performed C18 desalting, vacuum drying and buffer reconstitution. The samples were centrifuged prior to mass spectrometry analysis. Samples were analyzed by liquid chromatography-tandem mass spectrometry (LC-MS/MS) on a Q Exactive HF-X mass spectrometer (Thermo Scientific) coupled to a nano elute system. In total, 200 ng of sample was loaded onto a 25 cm × 75 μm column packed with 1.7 μm particles and equipped with IonOptics. Data were acquired in data-independent acquisition (DIA) mode via the diaPASEF method. False discovery rate (FDR) was set to 1.0% for both precursor ions and protein identification. All spectra were searched against the corresponding species database (mouse in this study) using trypsin digestion.

Carbamidomethylation (C, +57.02 Da) was defined as a fixed modification, and oxidation (M, +15.99 Da) was set as a variable modification. The details of peptides were list in the Supplementary Table S5.

### Cryo-EM sample preparation and data collection

Samples were buffer-exchanged to reduce glycerol from 10% to 2% while maintaining all other buffer components. To enrich low-concentration sperm proteasomes on grids, we used a simplified version of the IAAG strategy^51^. Graphene oxide-lacey carbon grids (300 mesh, EMR) were functionalized with 1 mM Pyr-NHS (Santa Cruz Biotechnology) in DMF for 1 hour at room temperature, washed three times with DMF, blotted dry, and air-dried for 10 min at room temperature. A 2 µL aliquot of sample was applied to grid and plunge-frozen in liquid ethane cooled by liquid nitrogen using a Vitrobot Mark IV (Thermo Scientific).

Cryo-EM data were collection using two transmission electron microscopes: a Titan Krios (Thermo Scientific) and a cryoARM300II (JEOL), both operated at 300 kV. For the Titan Krios transmission electron microscope, equipped with a Cs corrector and a K3 Summit direct electron detector (Gatan). Movies were recorded in super-resolution mode at a pixel size of 0.86 Å and a nominal magnification of 81,000×, resulting in a pexel size of 0.86 Å. Each movie was dose-fractioned into 40 frames, with a total accumulated dose of 44 e^-^/Å² on the specimen. Movies were collected using EPU software with defocus values ranging from –0.8 to –1.4μm. For the cryoARM300II transmission electron microscope (JEOL), equipped with a K3 Summit direct electron detector (Gatan). Movies were recorded in conted mode at a nominal magnification of 50,000 or 60,000, resulting in a pixel size of 1.07 or 0.91 Å. Each movie was dose-fractioned into 40 frames, with a total accumulated dose of 50 e^-^/Å² on the specimen. Movies were collected using SerialEM software with defocus values ranging from –0.8 to –1.5 μm.

### Cryo-EM 3D reconstructions

For mature sperm proteasome datasets, a total of 13,231 micrographs were processed using RELION 4.0^67^ and cryoSPARC 4.2.1^68^. Movies were aligned with MotionCorr2c^69^, CTF parameters were estimated with CTFFIND4^70^, and particles were autopicked with crYOLO 1.7.6^71^. After reference-free 2D classification, 652,019 good particles were retained. After heterogeneous refinement, two classes displaying better structural features (containing 399,009 and 167,546 particles, respectively) retained. Two rounds of no-align focused 3D classification resolved five proteasome classes: the 20S (201,413 particles), PAC1/2-20S-PAC1/2 (63,304 particles), PAC1/2-20S (99,764 particles), PA200-20S (101,250 particles), and PA200-20S-PAC1/2 (46,436 particles). After CTF refinement and Bayesian polishing, 20S and PAC1/2-20S-PAC1/2 particles were non-uniform refined in cryoSPARC with C2 symmetry to 2.76 Å and 2.88 Å resolution, respectively. PAC1/2-20S, PA200-20S, and PA200-20S-PAC1/2 particles were non-uniform refined in cryoSPARC with C1 symmetry to 3.01 Å, 3.12 Å, and 3.18 Å resolution, respectively. All the reported resolutions are based on the gold-standard FSC of 0.143 criterion.

For testis PA200-20S proteasome systems, a total of 11,520 micrographs were used for further processed. Particle picking with crYOLO and multiple rounds of 2D classification in cryoSPARC yielded 1,024,280 particles. Heterogeneous refinement in cryoSPARC resolved two major populations: PA200-20S (244,571 particles) and 20S proteasome (607,318 particles). Particles were further processed by no-alignment 3D classification, CTF refinement, and Bayesian polishing in RELION, followed by final non-uniform refinement in cryoSPARC. The final PA200-20S proteasome maps contained 146,669 and 582,942 particles and reached 2.97 Å resolution and 2.56 Å resolution, respectively, C2 symmetry was applied for the 20S map. The resolution was accessed based on the gold-standard criterion with Fourier shell correlation (FSC) at 0.143. All maps were sharpened using EMReady 2.3^72^, and figures were rendered using UCSF Chimera or UCSF ChimeraX^73^.

For human α4s-20S-PAC1/2 proteasome systems, a total of 13,347 micrographs were processed using cryoSPARC 4.7.1. After reference-free 2D classification, 2,504,487 good particles were retained. After heterogeneous refinement, two classes displaying better structural features (containing 403,353 and 997,787 particles, respectively) were retained. For 403,353 particles, several rounds of no-align focused 3D classification resolved four proteasome classes: PA200-20S (148,524 particles), PAC1/2-20S (94,487 particles), PA28-20S (46,985 particles), 20S (89,721 particles). For 997,787 particles, 3D classification resolved four proteasome assembly intermediates: PAC1-4/α-ring (192,094 particles), PAC1-2/α-ring/POMP/β2 (181,925 particles), Pre-13S (355,177 particles), and PAC1-2/α-ring/POMP/β2/β3/β1 (262,329 particles). After CTF refinement and reference based motion corretion, PA200-20S, PAC1/2-20S, PA28-20S, 20S, PAC1-4/α-ring, PAC1-2/α-ring/POMP/β2, Pre-13S and PAC1-2/α-ring/POMP/β2/β3/β1 particles were non-uniform refined in cryoSPARC with C1 symmetry to 2.97 Å, 3.01 Å, 3.14 Å, 3.02 Å, 3.53 Å, 3.47 Å, 3.29 Å and 3.18 Å resolution, respectively. Reported resolutions correspond to the gold-standardFSC of 0.143 criterion. All maps (except for PA28-20S) were sharpened using EMReady2. All figures were rendered using UCSF ChimeraX.

### Model building and validation

For initial templete, models of the bovine and human α4s subunits, bovine PA200, bovine PAC1-PAC2, and the bovine 19S portion of 26S proteasome were generated using AlphaFold3^74^ predictions, whereas the bovine or human 20S and PA200-20S proteasomes were based on previously published structures (listed in Supplementary Table S1-2). Initial models were docked into the corresponding cryo-EM density maps by rigid-body fitting in UCSF ChimeraX, and manually rebuilt in COOT^75^ to model features absent from the templates but resolved in our cryo-EM maps. The models were subsequently flexibly refined against the cryo-EM maps using ROSETTA^76^. Finally, the models were further refined against the cryo-EM maps using the phenix.real_space_refine module in Phenix^77^. All cryo-EM maps in study were listed in Supplementary Table S1-4.

## Data availability

The mass spectrometry proteomics data have been deposited in the ProteomeXchange Consortium (https://proteomecentral.proteomexchange.org) via the iProX partner repository under accession PXD078101. The dataset is currently kept private and available upon reasonable request from the corresponding author.

The cryo-EM maps determined for the proteasomes have been deposited at the Electron Microscopy Data Bank with the accession codes EMD-*****, EMD-***** and EMD-*****, and the associated atomic models have been deposited in the PDB with the accession codes ****, **** and ****, respectively.

## Acknowledgements

We thank the staff of the NCPSS Electron Microscopy, Database and Computing, and Protein Expression and Purification facilities for their instrument support and technical assistance. We also appreciate the technical support provided by the staff of the Integrated Laser Microscopy System (https://cstr.cn/31129.02.NFPS.CLMIS) at the National Facility for Protein Science in Shanghai (https://cstr.cn/31129.02.NFPS). Special thanks to Dr. Yifan Wang for setting up the protocol for purifying the PA200-20S from bovine spermatozoa.

This work was supported by grants from the National Key R&D Program of China (2024YFA1306204), Strategic Priority Research Program of CAS (XDB0570303), the NSFC (32570924 and 32130056), the National Key R&D Program of China (2024YFA1803102) and Shanghai Pilot Program for Basic Research from CAS (JCYJ-SHFY-2022-008).

## Author contributions

Y.C. initiated the project. Y.C., X.Z., L.S. and Z.H designed the experiements. L.S., X.Yuan., and C.C. prepared the endogenous PA200-20S proteasomes, 26S proteasomes from bovine mature spermatozoa and testis, collected cryo-EM data, and performed the structure reconstruction and data analysis. X.Z. generated the α4s-containing HEK293F cell lines and all related biochemical studies with assistance of X.L.. X.Z purified the PAC1/2-bound complexes from the engineered HEK293F cells; with Z.L. and C.C. collected related cryo-EM data with assistance of X.Ye, Z.L. and J. P. prefored recontructions. Proteomic profiling and data analysis were carried out by X.Z., L.S. with X.L.. Biochemical experiments were conducted by X.Z.. The 7G9 antibody was generated by T.W., X.Z, and T.C.. MS experiment was conducted by C.S. X.Z., Y.C., and L.S. wrote the manuscript with input from X.Yuan. and Z.H.

## Competing interests

All authors declare no competing interests.

## Supplementary Figures

**Fig. S1.**
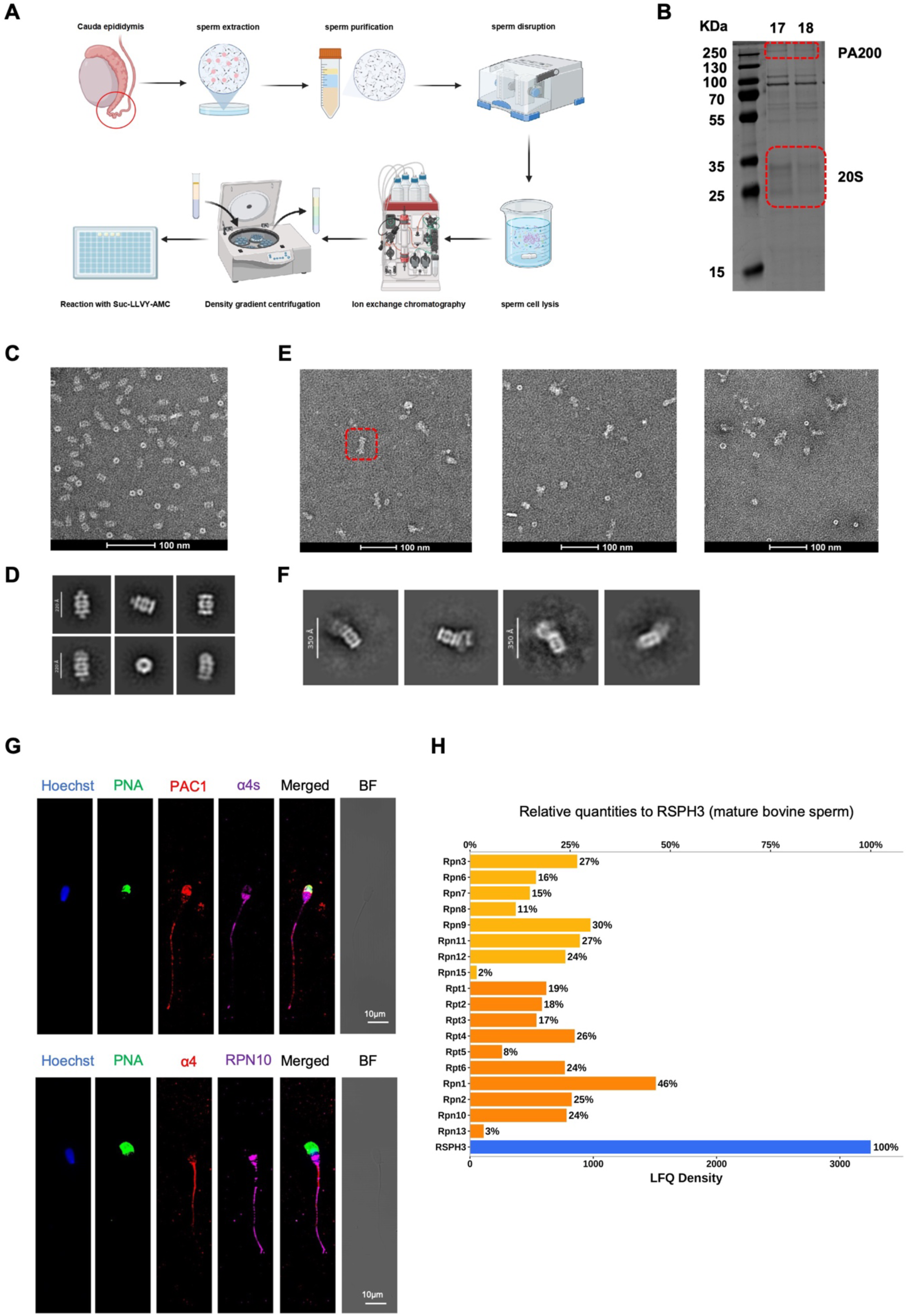
Purification and EM characterization of mature bovine sperm proteasomes. (A) Schematic workflow for proteasome purification from bovine spermatozoa, inclucing sequential cell lysis, differential centrifugation, ion-exchange chromatography, and glycerol-gradient centrifugation. (B) SDS-PAGE of purified sperm proteasomes, visualized by Coomassie Brilliant Blue staining. (C, D) Representative negative-stain (NS) EM micrographs (C) and reference-free 2D class averages (D) of purified proteasome samples. (E, F) Representative NS-EM micrographs (E) and reference-free 2D class averages (F) of the 26S proteasome fraction from mature bovine sperm, showing that 26S particles are only rarely observed. (G) Immunofluorescence showing PAC1, α4s, α4, and RPN10 localization in mature bovine spermatozoa. Hoechst (Hoechst33342) labels the nucleus; PNA marks the acrosome; BF, bright field. Scale bar: 10μm. (H) Relative abundance of 19S proteasome subunits in total mature bovine sperm lysate, normalized to the flagellar radial spoke head component RSPH3.

**Fig. S2.**
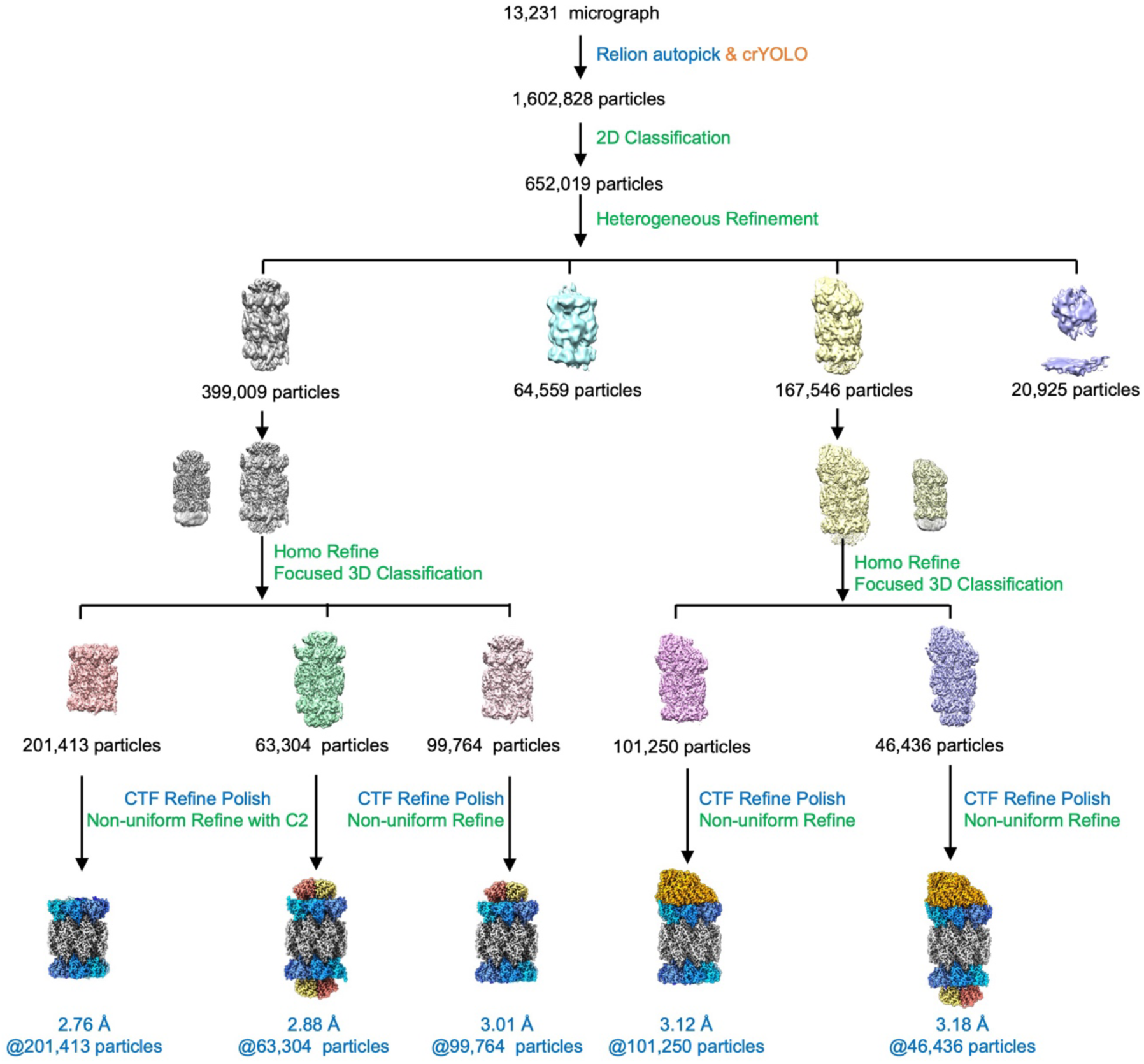
Cryo-EM data-processing workflow for mature sperm proteasome assemblies.

**Fig. S3.**
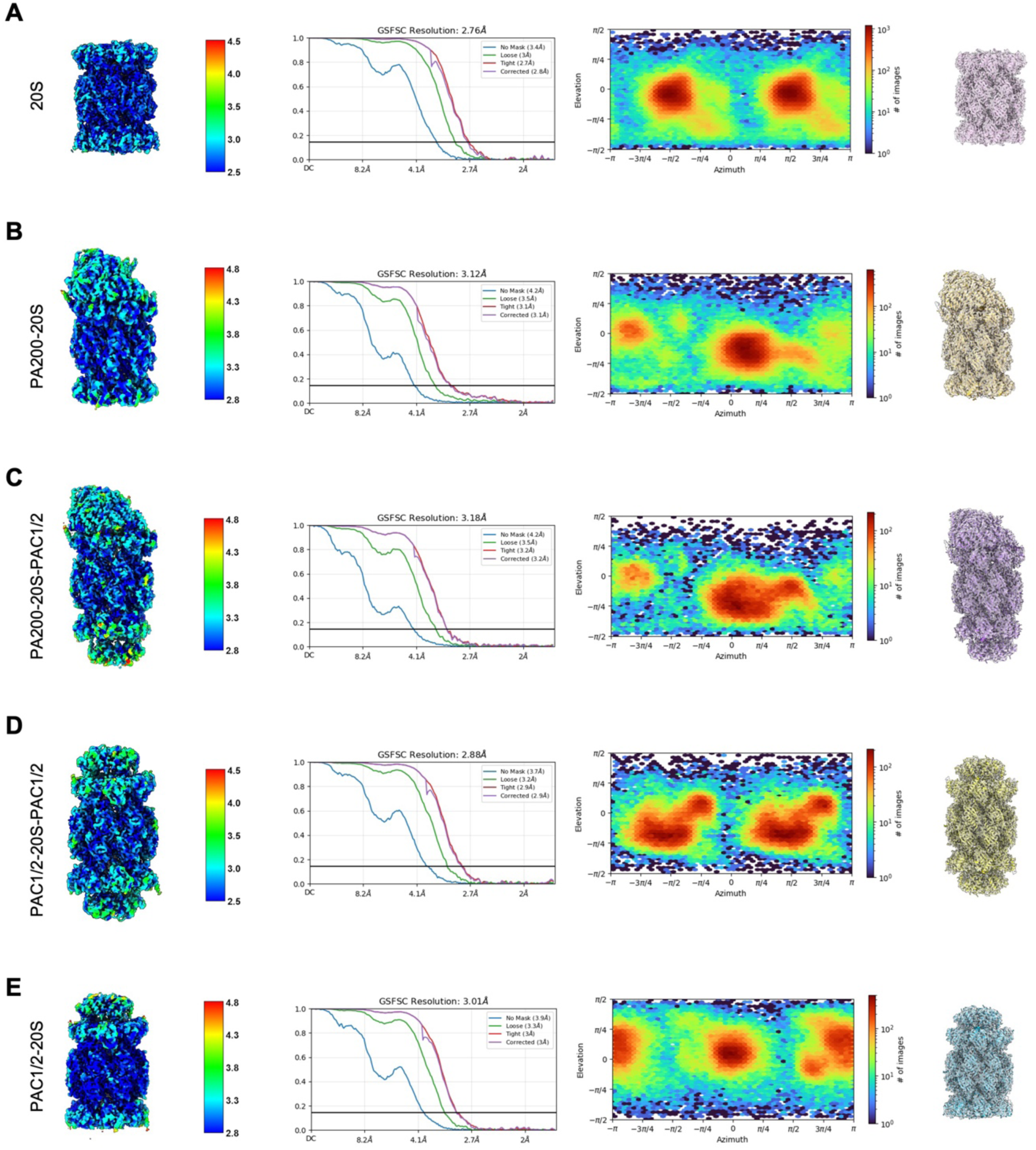
Resolution assessment and map-model fitting of mature sperm proteasome assemblies. (A-E) Local resolution estimates, Fourier shell correlation (FSC) curves, angular distribution of particle orientations, and map-model fitting for the free 20S (A), PA200-20S (B), PA200-20S-PAC1/2 (C), PAC1/2-20S-PAC1/2 (D), and PAC1/2-20S (E) complexes. FSC curves are plotted according to the gold-standard criterion.

**Fig. S4.**
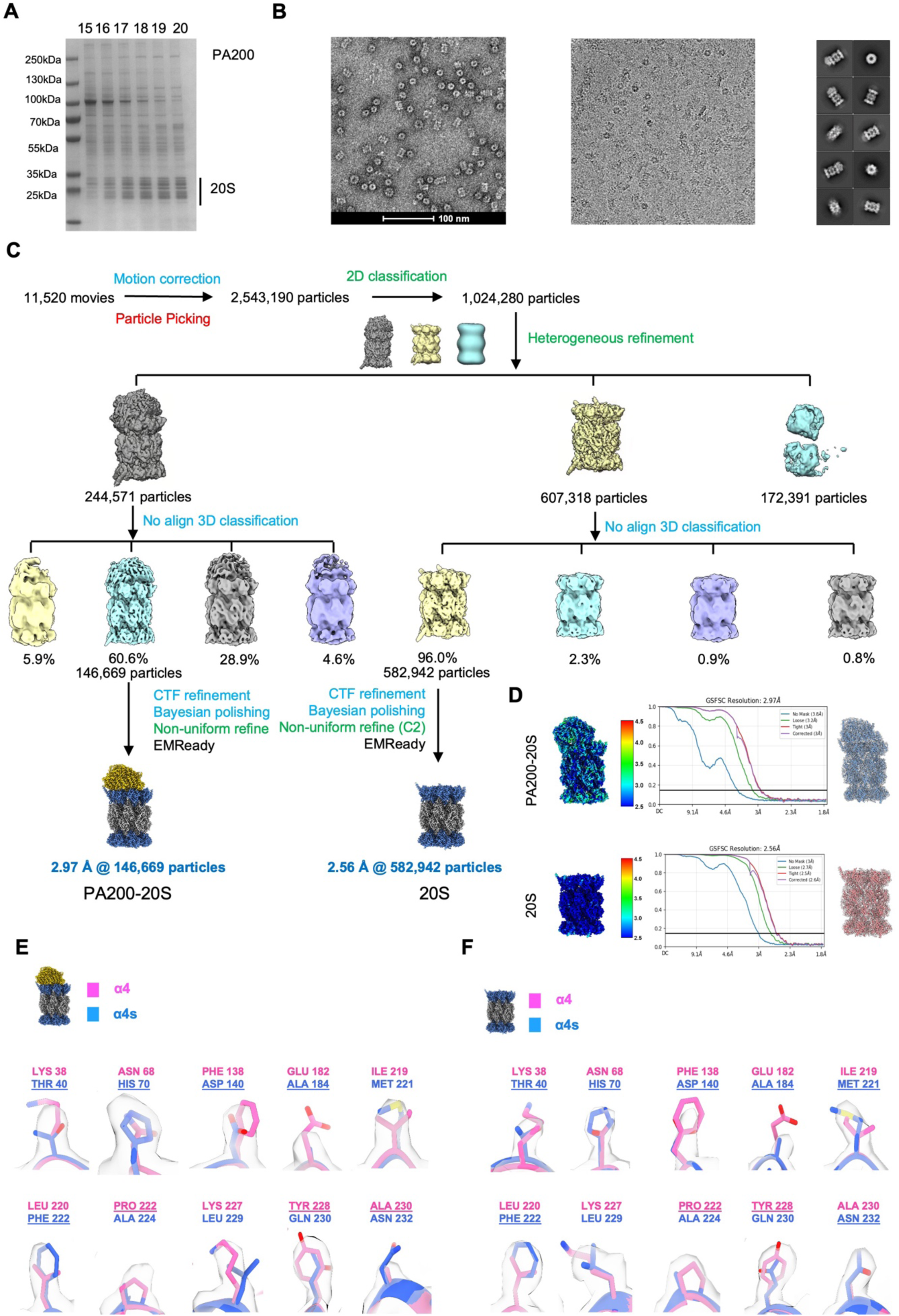
Purification and structural characterization of bovine testis proteasomes. (A) SDS-PAGE of purified testis PA200-20S proteasomes samples. (B) Representative NS-EM and cryo-EM micrographs, along with reference-free 2D class averages of purified testis PA200-20S proteasomes samples. (C) Cryo-EM data-processing workflow for the testis PA200-20S dataset. (D, E) Local resolution estimates, Fourier shell correlation (FSC) curves and representative map-model fitting for testis PA200-20S (D) and free 20S (E). FSC curves are plotted according to the gold-standard criterion. (E-F) Model-to-density comparisons at ten representative distinct α4/α4s positions in testis PA200-20S (E) and free 20S (F). The α4 model is shown in hot pink and α4s in royal blue; underlining residues denote the better-fitting model for each density.

**Fig. S5.**
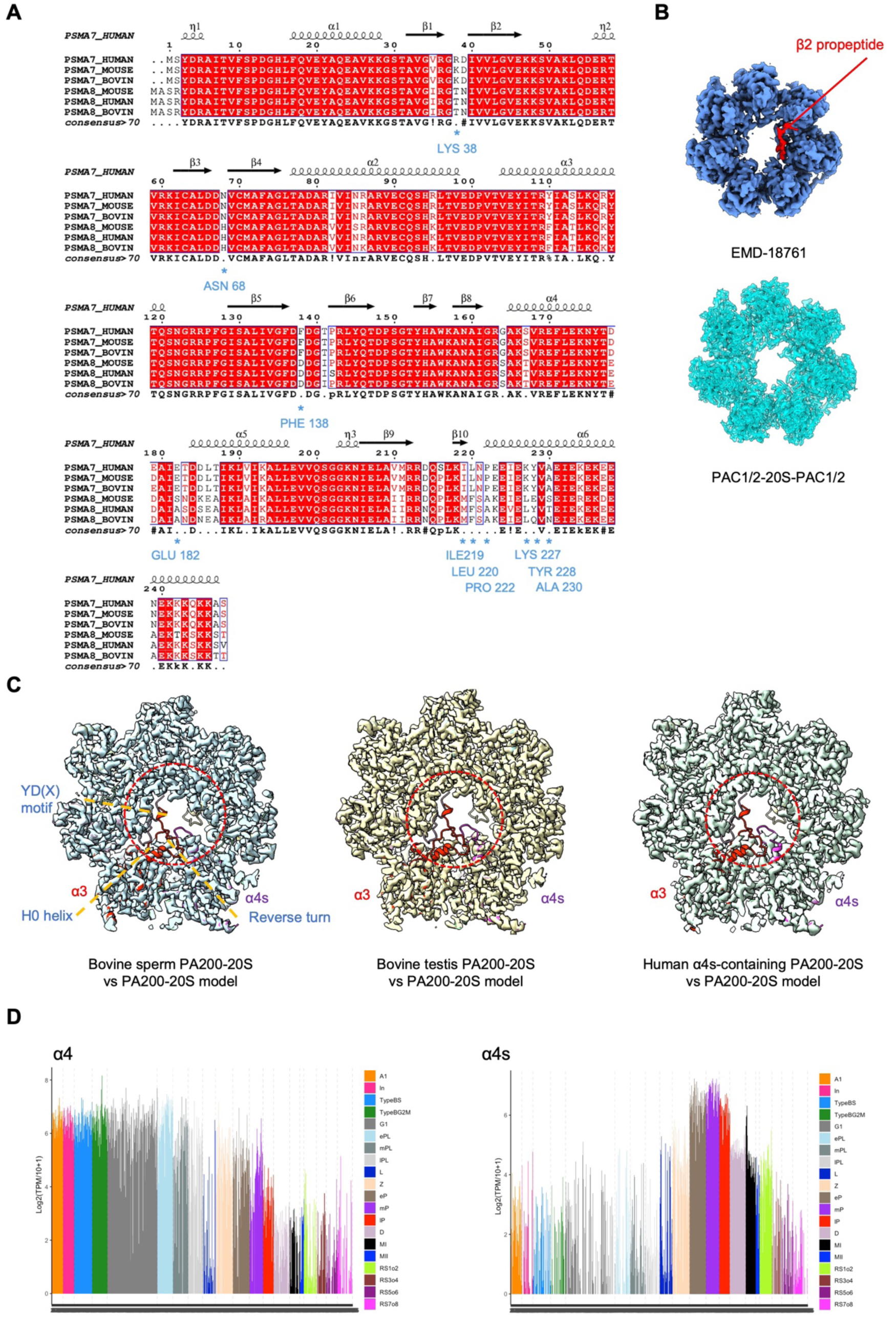
Sequence conservation and spermatogenic expression of mammalian proteasome α4 and α4s subunits. (A) Sequence alignment of α4 and α4s from human, mouse, and bovine. A consensus sequence is shown below the alignment. Diverge mostly in the C-terminal solvent-exposed regions, providing molecular markers (indicated by underneeth light blue stars) for distinguishing the two subunits in cryo-EM density maps. (B) Comparison of β2 propeptide density (red density) between the preholo 20S proteasome (EMDB: EMD-18761; upper panel) and the PAC1/2-20S-PAC1/2 complex (lower panel). (C) Structural comparison of the 20S gate in α4s-containing PA200-20S proteasomes across species and tissues, using our bovine sperm PA200-20S model as the reference/ruler. (D) Single-cell RNA sequencing (scRNA-seq) analysis of α4 and α4s expression dynamics during mouse spermatogenesis. Normalized gene expression levels are plotted across germ-cell stages, ranging from spermatogonia to post-meiotic round spermatids, with stage assignments indicated by color codes in the legend.

**Fig. S6.**
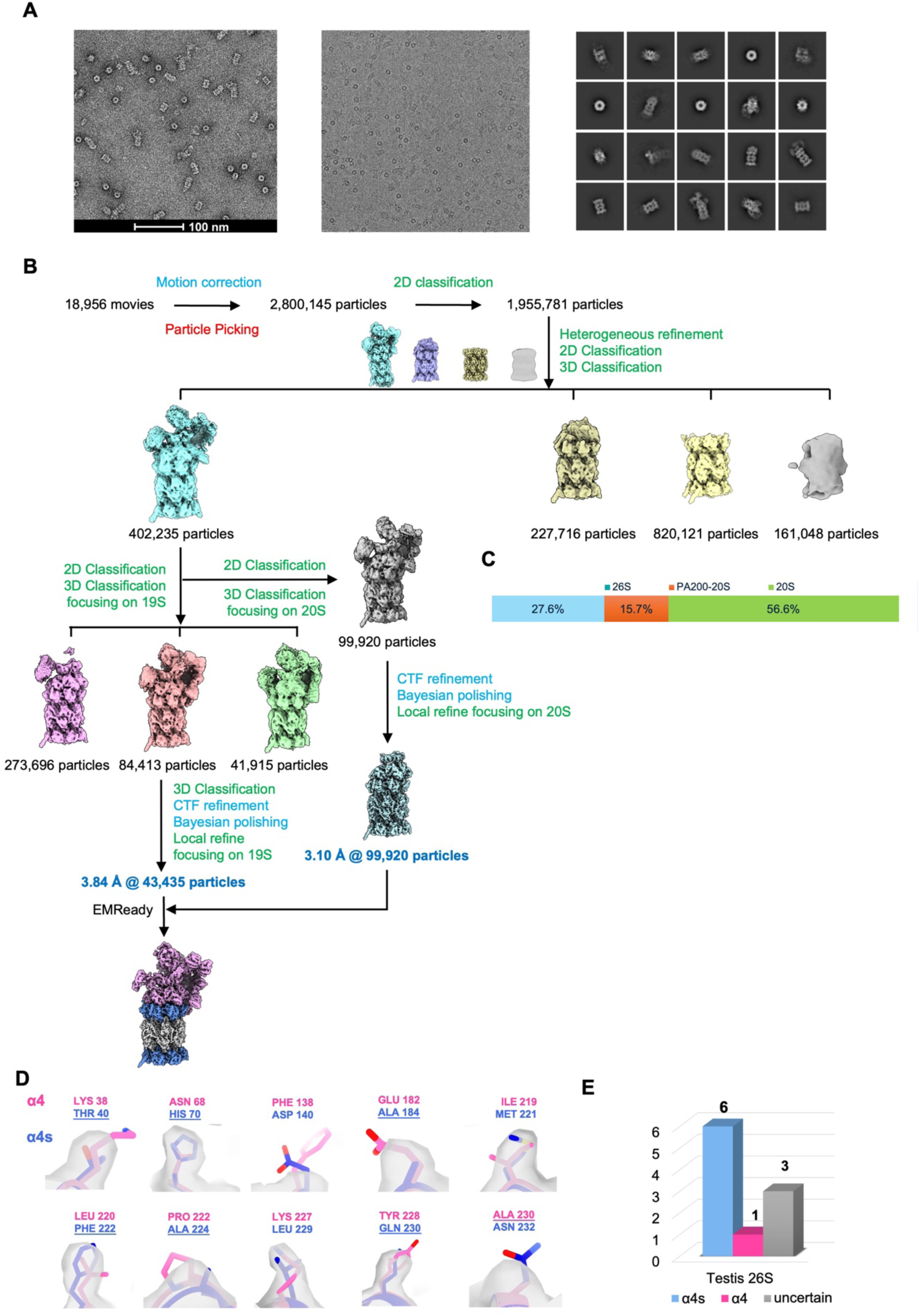
Cryo-EM data processing and structural characterization of the bovine testis 26S proteasome. (A) Representative NS-EM and cryo-EM micrographs, and reference-free 2D class averages of bovine testis 26S proteasome. (B) Cryo-EM data-processing workflow for the testis 26S proteasome dataset. (C) Relative abundance of 26S proteasome, PA200-20S, and 20S CP in bovine testis. (D) Model-to-density comparisons at ten representative distinct α4/α4s positions in bovine testis 26S proteasome. The α4 model is shown in hot pink and α4s in royal blue; underlined residues donote the better-fitting model for each density. (E) Quantification of α4 and α4s assignments based on diagnostic residue features.

**Fig. S7.**
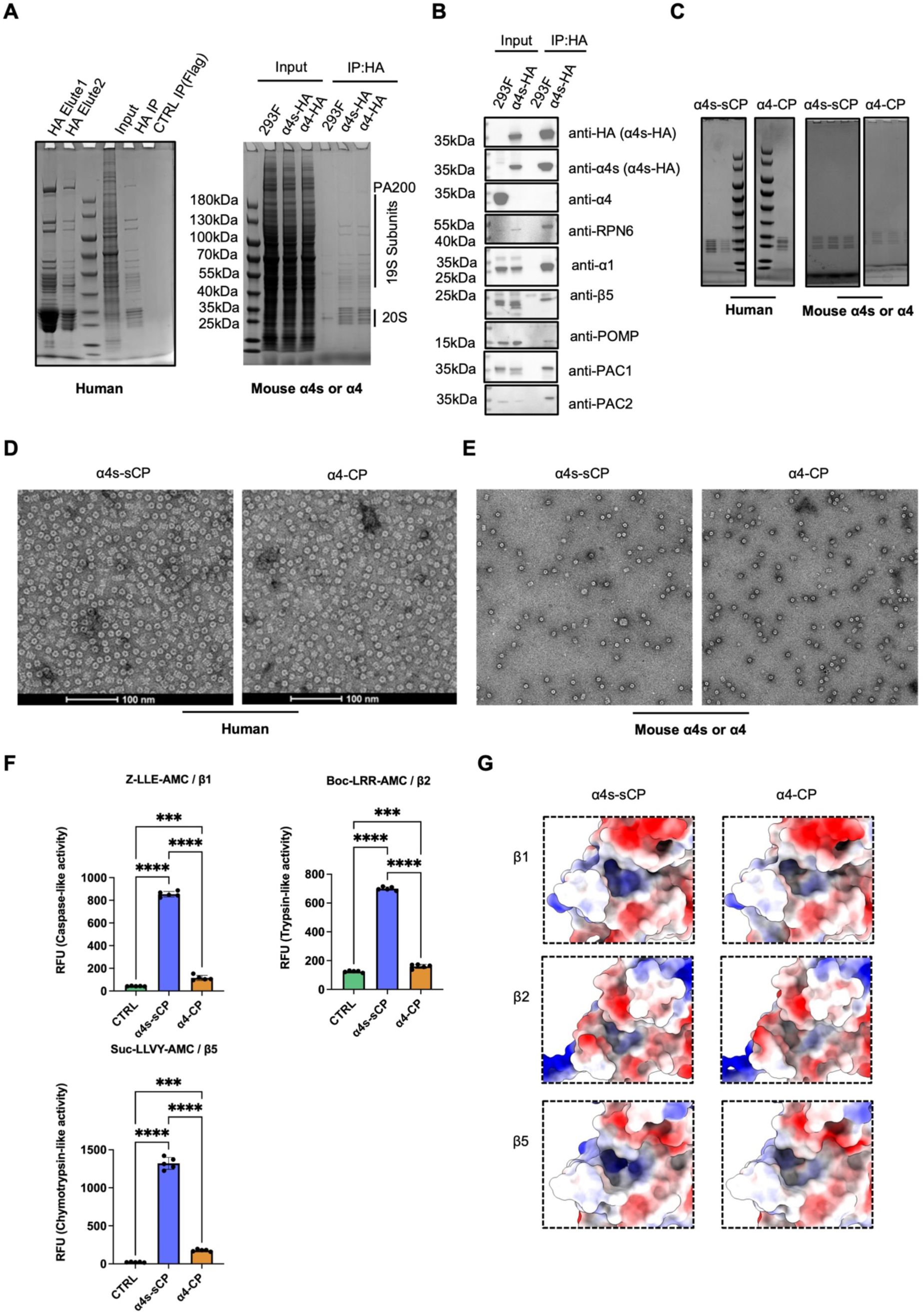
Biochemical characterization of α4s-containing proteasomes in α4-knockout HEK293F cells. (A) SDS-PAGE of HA affinity-purified eluates from α4-knockout HEK293F cells stably expressing human or mouse α4s-HA. (B) Co-IP and western blot validation of proteasome subunits and assembly chaperones in α4-knockout HEK293F cells stably expressing HA tagged mouse α4s. (C-E) SDS-PAGE (C) and NS-EM (D) of the human α4s-containing proteasome (α4s-sCP) or α4-containing 20S proteasomes (α4-CP), and NS-EM (E) of the moue α4s– or α4– containing 20S proteasomes, used for enzymatic activity assays. (F) Chymotrypsin-like, trypsin-like, and caspase-like proteasome activities of the purified mouse α4s-sCP or α4-CP. (G) Electrostatic potential surfaces of proteasome catalytic β subunits in human α4s-sCP and α4-CP.

**Fig. S8.**
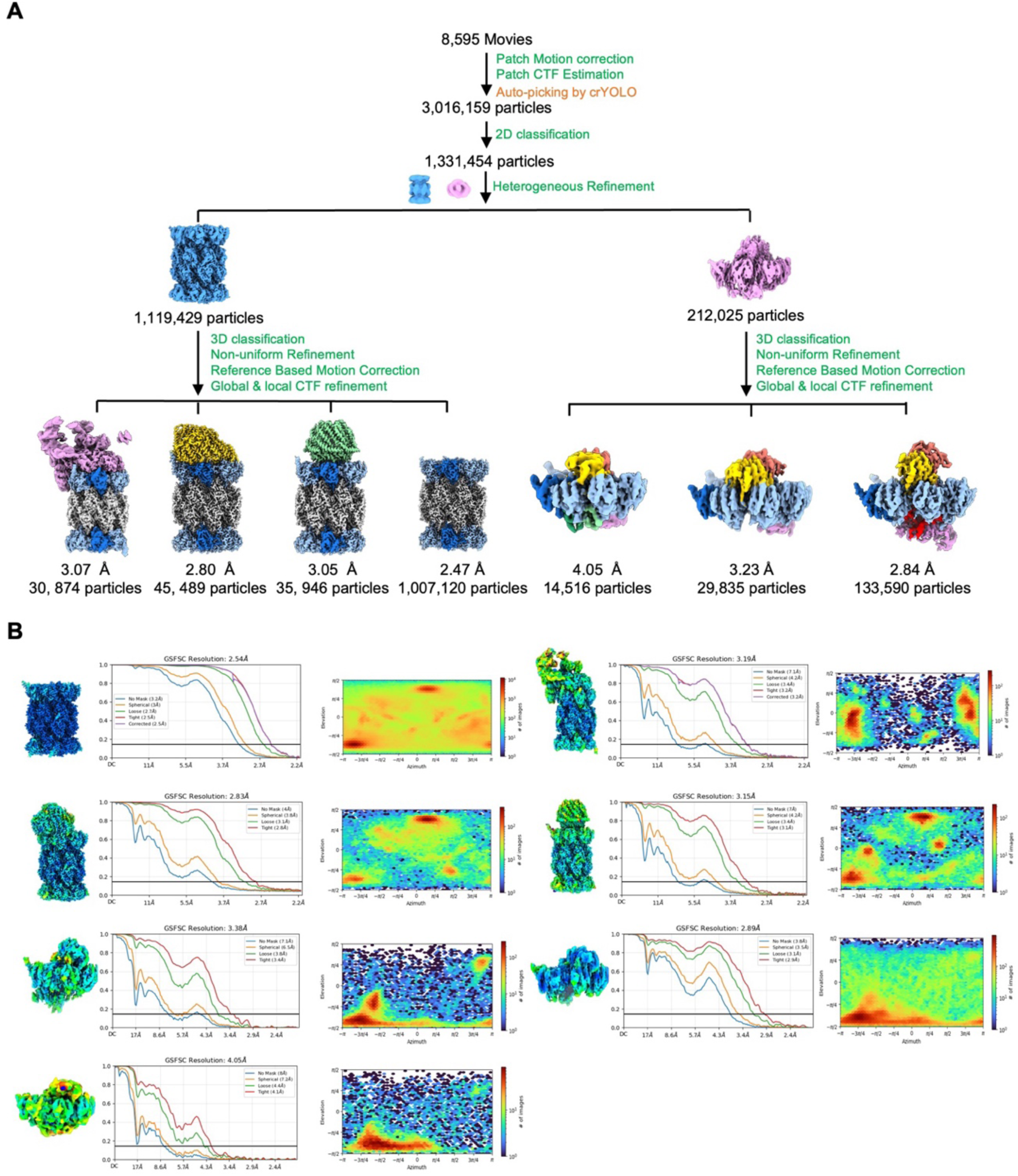
Cryo-EM processing of recombinant human α4s-containing proteasomes. (A) Cryo-EM data-processing workflow for human α4s-containing proteasomes. (B) Local resolution estimates, FSC curves, and map-model fitting for human α4s-containing proteasomes and assembly intermediates. FSC curves are shown according to the gold-standard criterion.

**Fig. S9.**
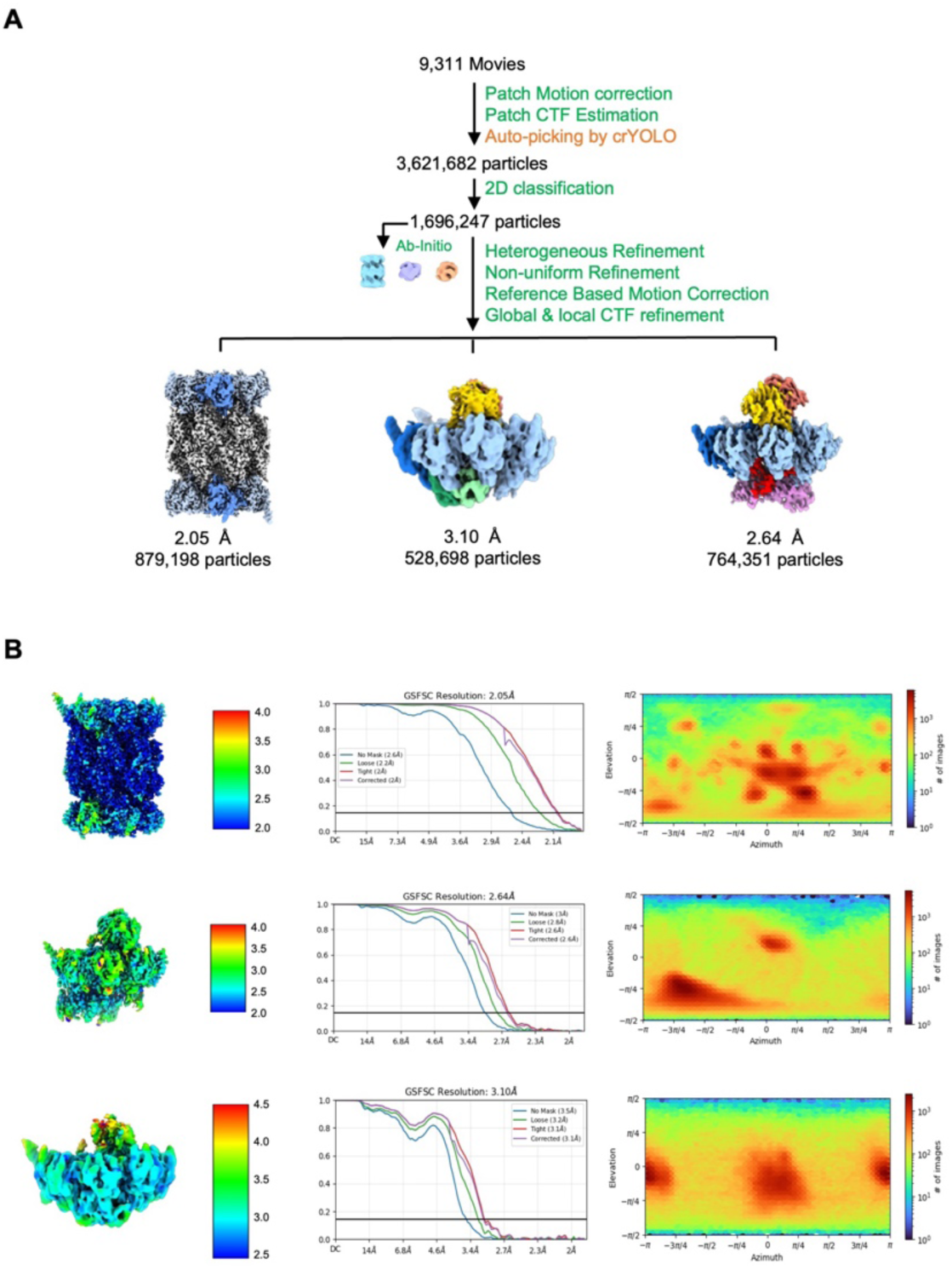
Cryo-EM processing of recombinant mouse α4s-containing proteasomes. (A) Cryo-EM data-processing workflow for mouse α4s-containing proteasomes. (B) Local resolution estimates, Fourier shell correlation (FSC) curves, and map-model fitting for mouse α4s-containing proteasomes and assembly intermediates. FSC curves are shown according to the gold-standard criterion.

**Fig. S10.**
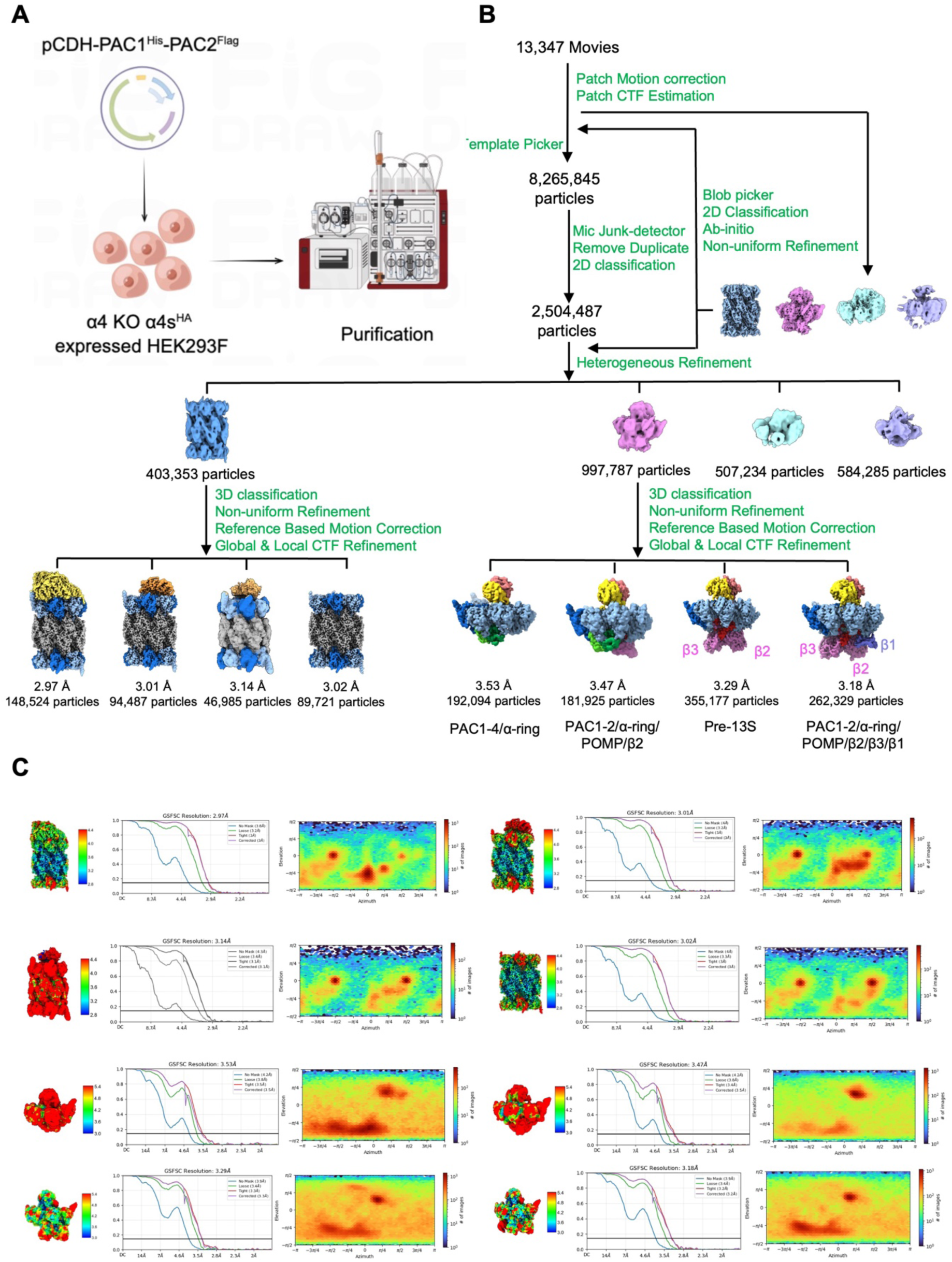
Cryo-EM processing workflow for the recombinant human PAC1/PAC2-bound α4s-containing proteasomes. (A) Schematic purification workflow for PAC1/PAC2-associated α4s-containing proteasomes from engineered HEK293F cells. (B) Cryo-EM processing workflow for recombinant human PAC1/PAC2-bound α4s-containing proteasomes. (C) Local resolution estimates, FSC curves, and map-model fitting for human α4s-containing proteasome complexes and assembly intermediates.

**Fig. S11.**
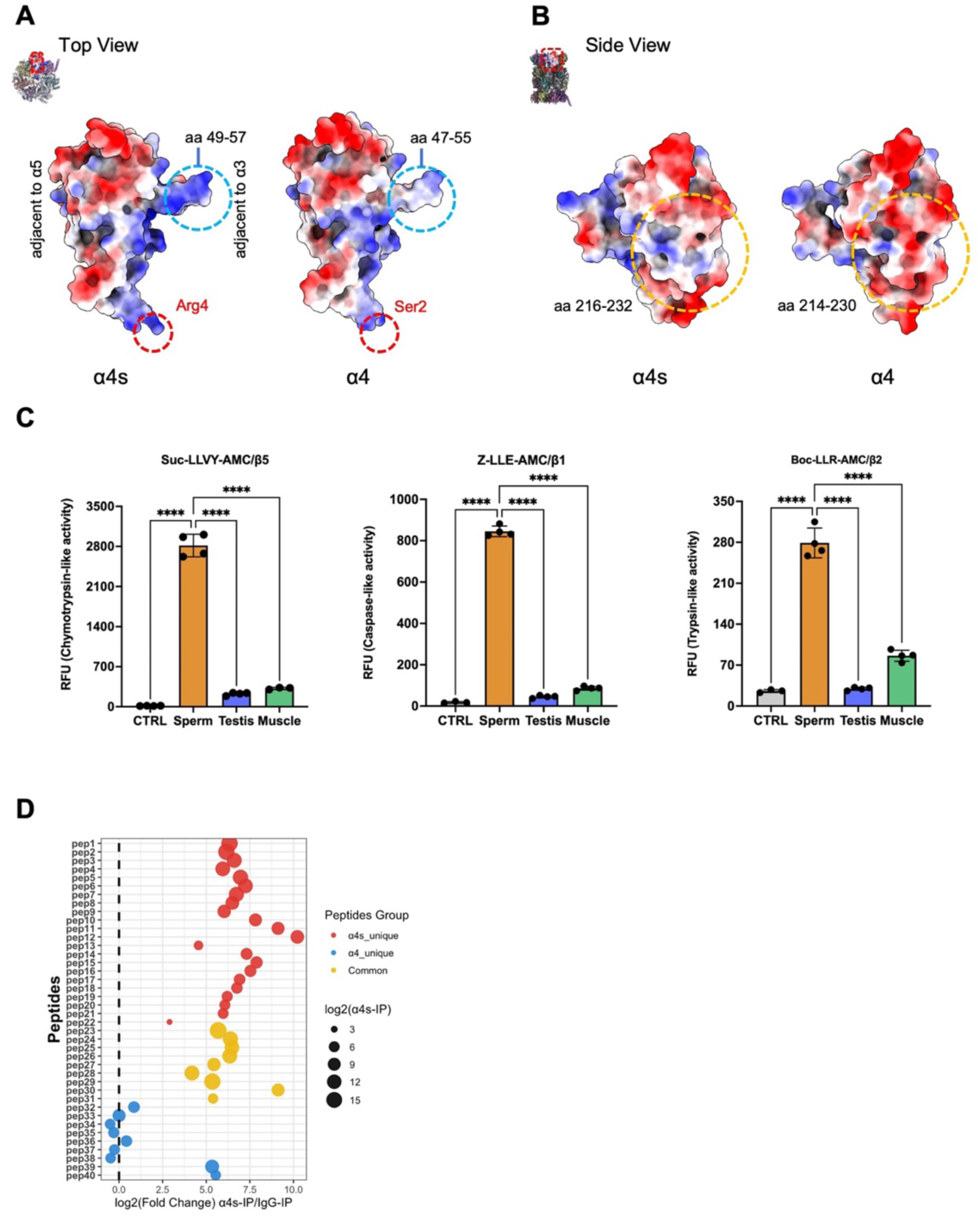
Functional and surface-property analysis. (A) Top-view electrostatic surface of the human α4s (left) and canonical α4 (right). Red dashed circles mark divergent charge patterns (α4s-Arg4 vs α4-Ser2); blue circles highlight the α3-adjacent divergent loop regions (residues 49–57 in α4s vs residues 47–55 in α4), which show increased positive surface potential in α4s. (B) Side-view electrostatic surfaces of human á4s-containing (left) and canonical á4-containing (right) 20S proteasomes. Yellow circles mark the divergent C-terminal loop region (residues 216–232) with distinct charge patterns (red: negative charge; blue: positive charge). (C) Proteasome activity assays for proteasomes in bovine mature sperm, testis, and muscle. (D) Enrichment of α4s and α4 peptides in mouse testicular sperm lysates via α4s immunoprecipitation/MS (Co-IP/MS). Peptide abundance (d.EG.TotalQuantity from co-IP/MS) was calculated as log_2_ fold change between α4s-IP and IgG-IP (x-axis). Peptides were categorized as α4s-unique (red), α4-unique (blue), or common (yellow). Dot size reflects log₂-transformed d.EG.TotalQuantity of the peptide in α4s-IP. The dashed line at log₂(FC)=0 marks the threshold for enrichment.

**Supplementary Table S1.** Summary statistics for cryo-EM data collection, processing and model validation of proteasomes from mature bovine spermatozoa.

|  | Mature Bovine Spermatozoa |  |  |  |  |
| --- | --- | --- | --- | --- | --- |
|  | PAC1-2/20S-PAC1/2 | PAC1-2/20S | PA200-20S/PAC1-2 | PA200-20S | 20S |
|  | (EMD-XXXXX) | (EMD-XXXXX) | (EMD-XXXXX) | (EMD-XXXXX) | (EMD-XXXXX) |
|  | (PDB-XXXX) | (PDB-XXXX) | (PDB-XXXX) | (PDB-XXXX) | (PDB-XXXX) |
| Data collection and processing |  |  |  |  |  |
| EM Equipment | Tian Krios |  |  |  |  |
| Detector | K3 |  |  |  |  |
| Energy filter slit width (eV) | 20 |  |  |  |  |
| Voltage (kV) | 81,000 |  |  |  |  |
| Electron exposure (e<sup>-</sup>/Å<sup>2</sup>) | 300 |  |  |  |  |
| Defocus range (μm) | 48 |  |  |  |  |
| Pixel size (Å) | -0.8~1.4 |  |  |  |  |
| Symmetry imposed | C2 | C1 | C1 | C1 | C2 |
| Initial particle images (no.) | 1602,828 | 1602,828 | 1602,828 | 1602,828 | 1602,828 |
| Final particle images (no.) | 63,304 | 99,764 | 46,436 | 101,250 | 201,413 |
| Map resolution (Å) | 2.88 | 3.01 | 3.18 | 3.12 | 2.76 |
| FSC threshold | 0.143 |  |  |  |  |
| Refinement |  |  |  |  |  |
| Initial model used | 1IRU | 1IRU | 6REY | 6REY | 1IRU |
| Model composition |  |  |  |  |  |
| Non-hydrogen atoms | 51,890 | 50,169 | 64,381 | 62,671 | 48,458 |
| Protein residues | 6,666 | 6,443 | 8,201 | 7,982 | 6,228 |
| B factors (Å<sup>2</sup>) |  |  |  |  |  |
| Protein | 48.62 | 54.24 | 64.12 | 73.84 | 41.60 |
| R.m.s. deviations |  |  |  |  |  |
| Bond lengths (Å) | 0.010 | 0.006 | 0.006 | 0.0068 | 0.006 |
| Bond angles (°) | 1.228 | 1.080 | 1.175 | 1.261 | 1.045 |
| Validation |  |  |  |  |  |
| MolProbity score | 1.82 | 1.70 | 1.74 | 1.91 | 1.55 |
| Clashscore | 3.92 | 3.51 | 3.60 | 4.38 | 3.30 |
| Poor rotamers (%) | 3.10 | 2.76 | 2.46 | 3.20 | 2.15 |
| Ramachandran plot |  |  |  |  |  |
| Favored (%) | 96.09 | 96.58 | 95.71 | 95.56 | 96.94 |
| Allowed (%) | 3.87 | 3.42 | 4.18 | 4.33 | 3.03 |
| Disallowed (%) | 0.05 | 0.00 | 0.11 | 0.11 | 0.03 |

**Supplementary Table S2.** Summary statistics for cryo-EM data collection, processing and model validation of proteasomes from bovine testis and human HEK293F cells.

|  | Bovine Testis |  | Human |
| --- | --- | --- | --- |
|  | 20S<br>(EMD-XXXXX)<br>(PDB-XXXX) | PA200-20S<br>(EMD-XXXXX)<br>(PDB-XXXX) | 20S<br>(EMD-XXXXX)<br>(PDB-XXXX) |
| Data collection and processing |  |  |  |
| EM Equipment | Tian Krios |  | cryoARM300 |
| Detector | K3 |  | K3 |
| Energy filter slit width (eV) | 20 |  | 20 |
| Voltage (kV) | 81,000 |  | 50,000 |
| Electron exposure (e<sup>-</sup>/Å<sup>2</sup>) | 300 |  | 300 |
| Defocus range (μm) | 50 |  | 50 |
| Pixel size (Å) | -0.8~1.4 |  | 1~2 |
| Symmetry imposed | C2 | C1 | C1 |
| Initial particle images (no.) | 2,543,190 | 2,543,190 | 3,016,159 |
| Final particle images (no.) | 582,942 | 146,669 | 1,007,120 |
| Map resolution (Å) | 2.56 | 2.97 | 2.47 |
| FSC threshold | 0.143 |  |  |
| Refinement |  |  |  |
| Initial model used | 1IRU | 6REY | 5VSF |
| Model composition |  |  |  |
| Non-hydrogen atoms | 48,732 | 63,020 | 48,871 |
| Protein residues | 6,258 | 8,026 | 6,280 |
| B factors (Å<sup>2</sup>) |  |  |  |
| Protein | 33.90 | 51.48 | 40.96 |
| R.m.s. deviations |  |  |  |
| Bond lengths (Å) | 0.0058 | 0.0064 | 0.005 |
| Bond angles (°) | 1.06 | 1.20 | 0.881 |
| Validation |  |  |  |
| MolProbity score | 1.39 | 1.76 | 1.14 |
| Clashscore | 2.79 | 4.11 | 3.28 |
| Poor rotamers (%) | 1.71 | 2.45 | 0.96 |
| Ramachandran plot |  |  |  |
| Favored (%) | 97.19 | 96.01 | 97.69 |
| Allowed (%) | 2.74 | 3.87 | 2.20 |
| Disallowed (%) | 0.06 | 0.13 | 0.11 |

**Supplementary Table S3.** Summary statistics for cryo-EM data collection and processing of proteasomes from human HEK293F cells expressing mouse or human α4s.

| | HEK293F cells expressing mouse $\alpha 4$s | | | HEK293F cells expressing human $\alpha 4$s | | | | | |
| --- | --- | --- | --- | --- | --- | --- | --- | --- | --- |
| | PAC1-4/α-ring | Pre-13S | 20S CP | PAC1-4/α-ring | PAC1-2/α-ring/POMP/$\beta 2$ | Pre-13S | PA200-20S | PA28-20S | 26S |
| Magnification | 60,000 |  |  | 50,000 |  |  |  |  |  |
| EM Equipment | JEOL cryoARM300 |  |  | JEOL cryoARM300 |  |  |  |  |  |
| Detector | Gatan K3 Camera |  |  | Gatan K3 Camera |  |  |  |  |  |
| Energy filter slit width (eV) | 20 |  |  | 20 |  |  |  |  |  |
| Voltage (kV) | 300 |  |  | 300 |  |  |  |  |  |
| Electron exposure ($e^-/\text{\AA}^2$) | 50 | | | 50 | | | | | |
| Defocus range ($\mu\text{m}$) | 0.6-1.8 | | | 0.6-1.8 | | | | | |
| Pixel size ($\text{\AA}$) | 0.91 | | | 1.07 | | | | | |
| Symmetry imposed | C1 | C1 | C1 | C1 | C1 | C1 | C1 | C1 | C1 |
| Initial particle images (no.) | 3,621,682 | 3,621,682 | 3,621,682 | 3,016,159 | 3,016,159 | 3,016,159 | 3,016,159 | 3,016,159 | 3,016,159 |
| Final particle images (no.) | 528,698 | 764,351 | 879,198 | 14,516 | 29,835 | 133,590 | 45,489 | 35,946 | 30,874 |
| Map resolution ($\text{\AA}$) | 3.10 | 2.64 | 2.05 | 4.05 | 3.23 | 2.84 | 2.80 | 3.05 | 3.07 |

**Supplementary Table S4.** Summary statistics for cryo-EM data collection and processing of proteasomes from human HEK293F cells expressing tagged human α4s, PAC1, and PAC2.

| | HEK293F cells with human $\alpha 4$s<sup>HA</sup> / PAC1<sup>His</sup>-PAC2<sup>Flag</sup> | | | | | | | |
| --- | --- | --- | --- | --- | --- | --- | --- | --- |
| | PAC1-4/α-ring | PAC1-2/$\alpha$ -<br>ring/POMP/β2 | Pre-13S | PAC1-2/α-<br>ring/POMP/β1/β2 | PA200-20S | PAC1/2-20S | PA28-20S | 20S CP |
| Magnification | 81,000 |  |  |  |  |  |  |  |
| EM Equipment | Titan Krios |  |  |  |  |  |  |  |
| Detector | Gatan K3 Camera |  |  |  |  |  |  |  |
| Energy filter slit width (eV) | 20 |  |  |  |  |  |  |  |
| Voltage (kV) | 300 |  |  |  |  |  |  |  |
| Electron exposure (e<sup>-</sup>/Å<sup>2</sup>) | 50 |  |  |  |  |  |  |  |
| Defocus range (μm) | 0.25~1.75 |  |  |  |  |  |  |  |
| Pixel size (Å) | 0.87 |  |  |  |  |  |  |  |
| Symmetry imposed | C1 |  |  |  |  |  |  |  |
| Initial particle images (no.) | 8,265,845 |  |  |  |  |  |  |  |
| Final particle images (no.) | 192,094 | 181,925 | 355,177 | 262,329 | 148,524 | 94,487 | 46,985 | 89,721 |
| Map resolution (Å) | 3.53 | 3.47 | 3.29 | 3.18 | 2.97 | 3.01 | 3.14 | 3.02 |

**Supplementary Table S5.** Antibodies and reagents used in this study.

| Names | Host | Cat No. | Supplier |
| --- | --- | --- | --- |
| 7G9 serum | Mouse | in house |  |
| 7G9 monoclonal antibody | humanization | in house |  |
| PSMA7 Rabbit mAb | Rabbit | A24916 | Company ABclonal, Inc. |
| PSMG1 Polyclonal antibody | Rabbit | 10335-1-AP | Proteintech Group, Inc. |
| PSMG2 Polyclonal antibody | Rabbit | 10959-1-AP | Proteintech Group, Inc. |
| PSME4 Polyclonal antibody | Rabbit | 18799-1-AP | Proteintech Group, Inc. |
| PSMB5 Rabbit pAb | Rabbit | A1975 | Company ABclonal, Inc. |
| PSMA6 Monoclonal antibody | Mouse | 67695-1-Ig | Proteintech Group, Inc. |
| Anti-PSMD14 antibody | Rabbit | EPR4257 | Abcam |
| PSMD4 Monoclonal antibody | Mouse | 66179-1-Ig | Proteintech Group, Inc. |
| clone M2, affinity isolated antibody | Mouse | F1804 | Sigma |
| Anti-mouse IgG, HRP-linked Antibody |  | #7076 | Cell Signaling Technology, Inc. |
| Goat Anti-Rabbit IgG H&L(HRP) |  | ab97051 | Abcam |
| goat anti-Rabbit IgG Alexa Fluor™ 555 |  | A-21428 | Thermo Fisher Scientific |
| AF647-labeled goat anti-mouse IgG (H+L) |  | A0473 | Beyotime |
| Peanut agglutinin |  | L32458 | Thermo Fisher Scientific |
| antifade mounting medium |  | P0133 | Beyotime |
| FuturePAGETM 4-20% 15 Wells |  | ET15420LGel | ACE |
| Anti-DYKDDDDK Affinity Beads |  | SA042100 | Smart-Lifesciences |
| Anti-HA Affinity Beads |  | SA068100 | Smart-Lifesciences |
| cOmplete EDTA-free PIC |  | 04693132001 | Roche Diagnostics GmbH |
| Western BLot Ultra Sensitive HRP |  | T7104A | Takara |
| Lipo8000 |  | C0533 | Beyotime |
| 4% Tissue Fix Solution |  | T17024 | Saint-Bio Biotech |
| immunostaining permeabilization buffer |  | M20254 | Saint-Bio Biotech |

**Supplementary Table S6.**
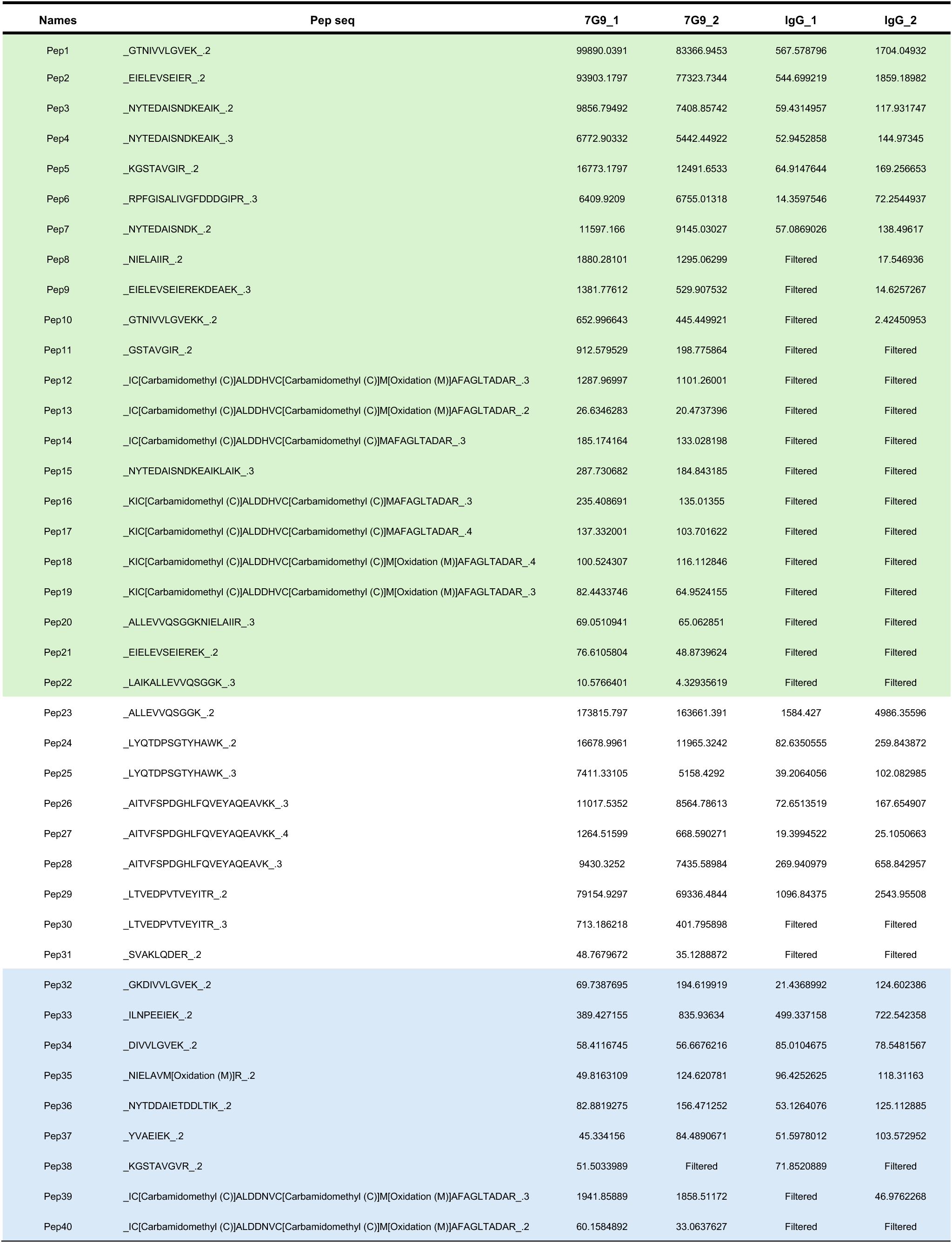
α4s and α4 peptides in mouse testicular sperm lysates via α4s immunoprecipitation/MS (Co-IP/MS)

## Notes

### Competing Interest Statement

The authors have declared no competing interest.

